# Age-associated spontaneous activation of BMP signaling inhibits the canonical Wnt pathway through ICAT to promote osteoarthritis

**DOI:** 10.1101/2024.10.02.616036

**Authors:** Bhupendra Kumar, Sayeda Fauzia Iqbal, Ankita Jena, Shuchi Arora, Saahiba Thaleshwari, Makoto Mark Taketo, Amitabha Bandyopadhyay

## Abstract

Osteoarthritis (OA) is a highly prevalent and debilitating musculoskeletal disorder that affects billions of aging individuals worldwide, causing chronic pain, impaired mobility, and a substantial decline in quality of life, yet effective disease-modifying therapies remain unavailable. Moreover, existing experimental models, however, inadequately capture the complex, age-related factors that contribute to OA pathogenesis, limiting our understanding of underlying molecular mechanisms and hindering therapeutic development. To address this gap, we examined mouse articular cartilage (AC) during natural aging, without surgical or chemical intervention. In young AC, BMP ligand expression is restricted to the bone–cartilage junction, limiting BMP signaling despite widespread BMPR1A receptor expression, while Wnt/β-catenin signaling predominates in the outer layers. With aging, BMP ligand expression expands throughout the cartilage, leading to widespread activation of BMP signaling, which induces ICAT expression, a known inhibitor of Wnt/β-catenin signaling. This elevated BMP signaling triggers chondrocyte hypertrophy while concurrently suppressing Wnt/β-catenin activity. Moreover, genetic or pharmacological activation of BMP signaling in young AC recapitulates the cellular and molecular features of aged cartilage, independent of chronological age. Collectively, our findings demonstrate that an age-dependent shift in BMP-Wnt signaling disrupts AC homeostasis and drives OA progression in mice.

**Author Summary:** Despite the prevalence of osteoarthritis in aging individuals, there are no effective treatments that can stop or reverse disease progression. In this study, we investigated how joint cartilage changes during natural aging in mice. We observed that healthy, young articular cartilage exhibits robust Wnt/β-catenin with minimal BMP siganling. With age, this balance shifts, with BMP signaling becoming dominant, coupled with reduced Wnt/β-catenin activity. This disruption of Wnt-BMP homeostasis induces the pathogenesis of OA. Importantly, local inhibition of BMP signaling significantly alleviates the severity of the disease. Overall, these results show that age-associated changes in key signaling pathways drive OA progression and suggest potential targets for therapeutic intervention.

**Graphical abstract:** 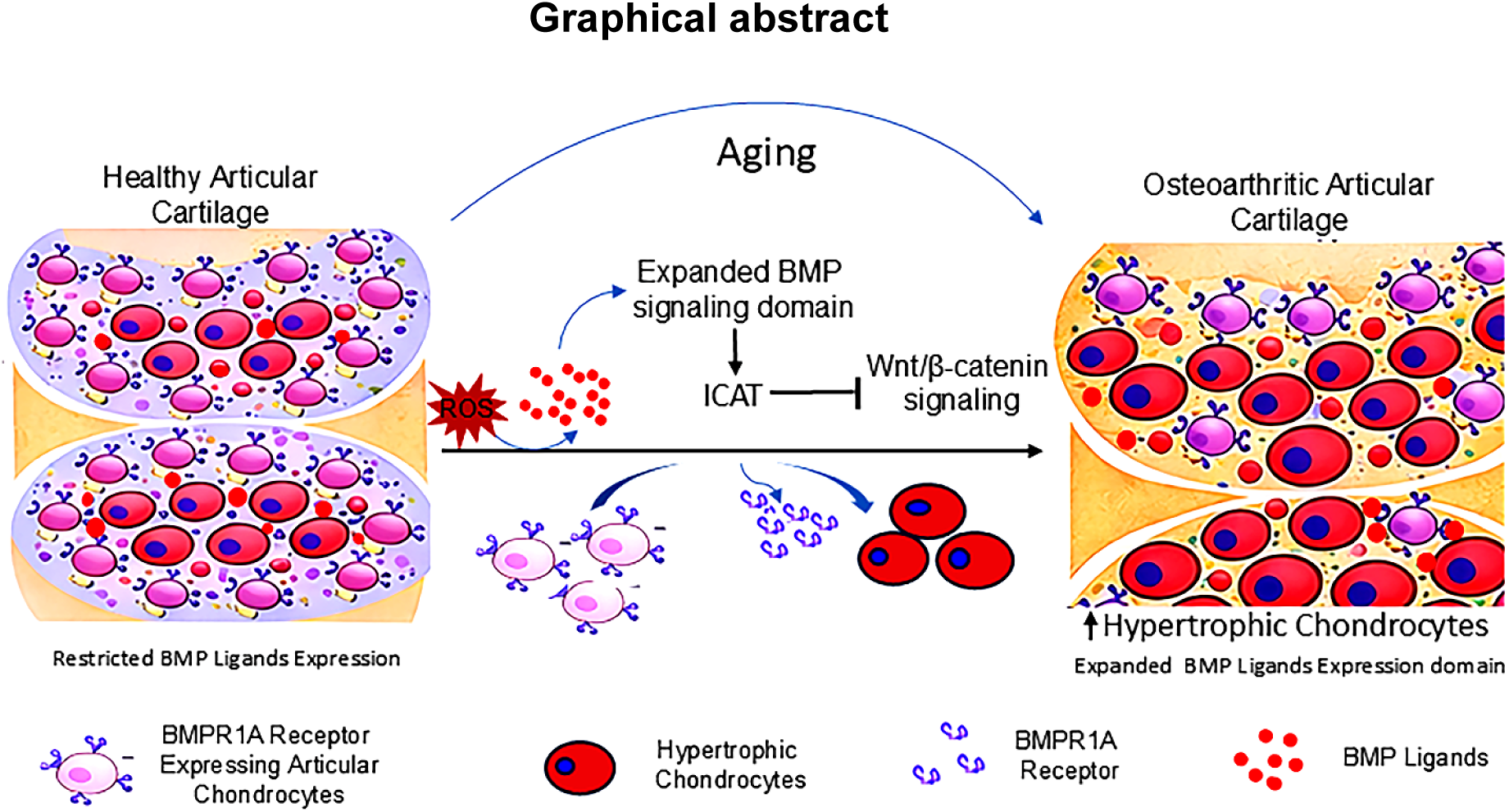

## Introduction

Osteoarthritis (OA) is one of the most common musculoskeletal disorders that affects the synovial joints, primarily in the elderly population[1,2]. It is a medical condition exhibiting one of the fastest growth rates in terms of <u>Y</u>ears <u>L</u>ived with <u>D</u>isability (YLDs) after mental and behavioural disorders[3]. Its progression manifests as gradual changes in the structure and composition of articular cartilage (AC)[4]. Briefly, cells of the AC undergo chondrocyte hypertrophy, aberrant calcification, and osteophyte formation with low-grade inflammation, failing the joint’s functionality[5]. Interestingly, the pathophysiology of OA and embryonic endochondral bone formation share many molecular similarities[5]. Similar to endochondral ossification during OA progression, healthy type II collagen (Col II)-expressing cartilage cells undergo hypertrophy by expressing type X collagen (Col X) and MMP 13. These hypertrophic chondrocytes recruit blood vessels by expressing VEGF and differentiate into osteoblasts that secrete calcified matrix rich in collagen type 1 (Col I)[5–8].

Old age is the most common predisposing factor for OA[9]. Yet, investigations into the molecular pathology of OA are predominantly conducted using PTOA(Post-traumatic osteoarthritis) or chemically induced models, where the animals used are almost always young, that is, before they become naturally prone to develop OA[10–13]. These precocious, induced models likely fail to capture the complex, multifactorial processes underlying the natural onset of OA in the aging population[9,14]. Also, the available literature on OA patients, for obvious reasons, is limited to characterisation of the disease pathology in its terminal stages rather than elucidating the molecular pathways involved in the onset and/or progression of the disease[15,16]. In this context, it should be noted that Rowe *et. al*. demonstrated that the factors involved in age-associated OA are likely to be different from the ones involved in the pathogenesis of PTOA[17] This lack of detail presents a significant challenge for the scientific community in developing a molecular-level understanding of age-associated OA [10,18].

BMP signaling is known to be critically needed for endochondral ossification. The mice lacking *bmp2* and *bmp4* genes, fail to develop endochondral bone[19]. Further, conditional limb-specific BMP signaling deprivation causes delayed chondrocyte hypertrophy and thereby endochondral ossification[20]. Additionally, intra-articular injection of the BMP signaling inhibitor, noggin, reduces the severity of PTOA in rats[21]. Further, in vitro-produced cartilage retains its native phenotype when treated with a BMP signaling inhibitor before implantation in *ex vivo* animal models [22]. Moreover, suppression of BMP signaling also prevents chondrocyte hypertrophy in micro-cartilage models of clinical OA [23]. Jaswal *et. al.* demonstrated that genetic or pharmacological inhibition of BMP signaling mitigates surgically induced OA in mice [24,25]. However, as highlighted by Lyons and Rosen, it is still unclear whether activation of BMP signaling is linked to age-related osteoarthritis[25].

In 2015, Ray et.*al*. demonstrated that strict spatial restriction of BMP and Wnt signaling domains during early embryonic development is critical for simultaneous differentiation of transient ( bone precursor) and permanent (joint or articular) cartilage [26]. Studies demonstrated that, on the one hand, the canonical Wnt pathway is essential for the development of AC [27,28], yet this pathway also promotes hypertrophic differentiation of chondrocytes[29]. It is also puzzling that both loss- and gain-of-function of canonical Wnt signaling in adult AC induce OA-like degenerative changes[30–32]. Further, elevated levels of Wnt ligands have been linked to OA, and their inhibition has been shown to confer protective effects in PTOA of mice[33]. Therefore, the precise role of the canonical Wnt pathway to adult AC homeostasis remains unclear, particularly in the context of age-related cartilage degeneration [34,35].

In this study, we found that age-associated expansion of the BMP-ligand expression domain elevates BMP signaling throughout the murine AC. This increased BMP activity induces ICAT, which suppresses canonical Wnt/β-catenin signaling and drives degenerative OA-like changes. Moreover, transient or genetic upregulation of BMP signaling was sufficient to recapitulate the full spectrum of cellular and molecular features of OA in mice, regardless of their chronological age. Our data suggest that a shift from canonical Wnt to BMP-dominant signaling in aging AC represents a key degenerative mechanism driving age-associated OA.

## Results

### 1. OA progression accompanies with increased BMP signaling in aging AC

The primary aim was to systematically evaluate the changes in the AC across stages from juvenile to aged mice, and to determine the earliest time points at which degenerative changes arise during aging. We used C57BL/6J wild-type mice without genetic or pharmacological intervention and examined cellular and molecular changes. To evaluate these changes, knee joints were collected from mice at 6, 18, 24, 30, and 36 months of age for the analysis. Articular chondrocytes normally express Col II, but its expression declines during OA progression, accompanied by an increase in the hypertrophic marker Col X [36,37]. We therefore evaluated the expression of Col II and Col X in AC across different age groups of mice. We found abundant Col II in the 6-month-old AC samples. The immunoreactivity for Col II steadily declined between 18 and 36 months of age (Fig.1B-B′′′′ and Fig.1G). Moreover, immunoreactivity for Col X showed a marked increase, indicating enhanced chondrocyte hypertrophy beginning at 18 months of age, which peaked around 36 months (Fig.1C-C′′′′, and Fig.1H). Furthermore, to assess the overall integrity of AC, we performed safranin O/fast Green staining, a method traditionally used for evaluating AC[38]. Intense Safranin O staining in 6-month-old mice AC reflects abundant proteoglycan content, which gradually declined from 18 months onward, reaching its lowest levels at 30 and 36 months of age (Fig.1D-D′′′′,1E-E′′′′). The severity of OA in murine AC was assessed using Safranin O-stained sections from 6-month-old to 36-month-old mice, following OARSI histopathological guidelines[39]. Significantly higher OARSI scores at 24, 30, and 36 months compared to 6-month-old mice highlight a clear link between OA progression and aging (Fig.1I).

**Fig. 1.**
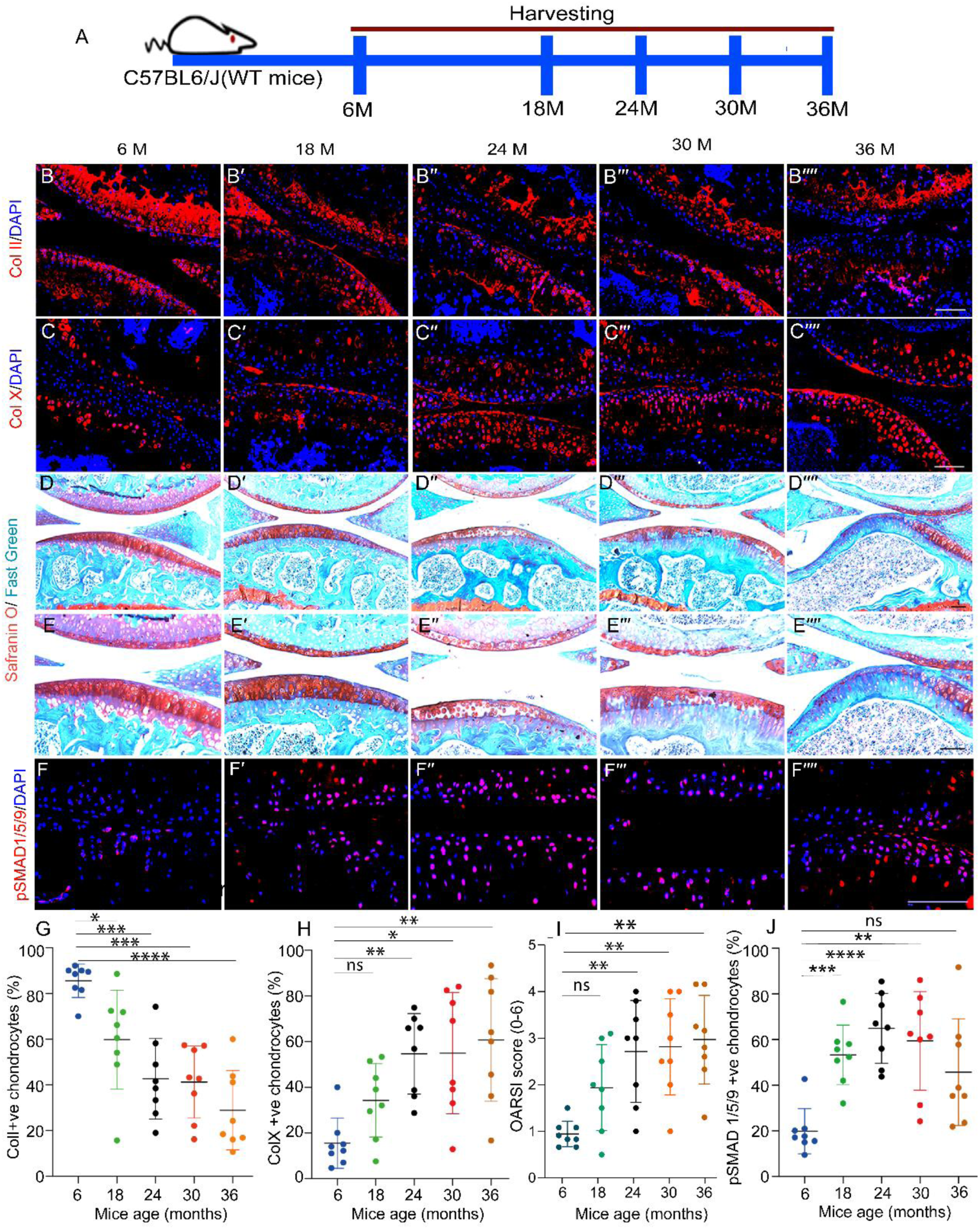
Age-associated OA etiology associated with up-regulated BMP signaling. (A) Schematic representation depicting selected experimental time points for analysis of molecular signatures of aging C57BL/6J WT mice. (B-F′′′′) Longitudinal sections through the adult knee joints of C57BL/6J (WT mice) harvested at 6 months (B-F), 18 months (B′-F′), 24 months (B′′-F′′), 30 months (B′′′-F′′′) and 36 months (B′′′′-F′′′′) of age. Immunoreactivity for ColII (B-B′′′′), Col X (C-C′′′′), Safranin O staining (D-D′′′′), Safranin O staining inset (E-E′′′′) (Scale bar: 20µm). (F-F′′′′) Immunoreactivity for pSMAD 1/5/9. (G) Quantification data for ColII, Brown-Forsythe, and Welch ANOVA were performed along the five sets, and p < 0.0001(****). We compared the means of 6 months vs. 18 months; p = 0.0361 (*), 6 months vs. 24 months; p = 0.0004 (***), 6 months vs. 30 months p = 0.0001 (***), and 6 months vs. 36 months p <0.0001 (****). (H) Quantification data of Col X, Brown-Forsythe, and Welch ANOVA were performed along the five sets, and p < 0.0001 (****). We compared the means of 6 months vs. 18 months; p = 0.0569 (ns), 6 months vs. 24 months; p = 0.0007 (***), 6 months vs. 30 months, p = 0.0115 (*), and 6 months vs. 36 months, p =0.0052 (**). (I) OARSI score, Brown-Forsythe, and Welch ANOVA were performed along the five sets; we compared 6 months vs 18 months; p = 0.0562 (ns), 6 months vs 24 months; p = 0.0073(**), 6 months vs 30 months; p=0.0037(**), 6 months vs 36 months; p=0.0013(**). (J) Quantification data of pSMAD1/5/9, Brown-Forsythe, and Welch ANOVA were performed along the five sets, and p < 0.0001 (****). We compared the means of 6 months vs. 18 months; p = 0.0002 (***), 6 months vs. 24 months; p < 0.0001 (****), 6 months vs. 30 months, p = 0.0029 (**) and 6 months vs. 36 months, p <0.0534 (ns). Scale bar = 100 µm; n=8 per group. Error bars represent the mean ± standard deviation (S.D.).

An increased vascularisation and ectopic bone formation (osteophytes) in the AC is the prominent hallmark of OA progression[40,41]. To assess vascularisation and osteophyte formation, we performed micro-CT imaging to detect the presence of blood vessels and osteophytes in the AC of 6-month-old and 30-month-old mice. Samples from the 30-month-old group (Fig. S1A′-1B′) exhibited a pronounced increase in vascularisation (shown as intensified red color), compared to samples from the 6-month-old group (Fig. S1A-1B). Further, cross-sectional micro-CT images revealed that bone marrow blood vessels extended to and covered the articular surface in 30-month-old samples (Fig.S1B′), a feature that was absent in 6-month-old mice. (Fig.S1B). Additionally, 30-month-old mice AC showed signs of excessive mineralisation and osteophyte formation, which were absent in 6-month-old mice AC (Fig.S1A and S1B, Fig.S1A′ and S1B′). The above data strongly suggest that C57BL/6J mice exhibit molecular and cellular features of OA progression. Jaswal *et al.* have identified a critical role for BMP signaling in the pathogenesis of PTOA[24]. However, given the multifactorial etiology of OA, the molecular drivers may differ across disease contexts. We therefore examined the status of BMP signaling in aging AC. To this end, we performed immunohistochemistry for pSMAD1/5/9, a readout of BMP signaling, across all examined time points. Minimal immunoreactivity was observed at 6 months of age, followed by a progressive increase thereafter, with a subsequent reduction at 36 months (Fig.1F-F′′′′ and Fig.1J). Collectively, these results demonstrate an age-dependent upregulation of BMP signaling in AC, closely associated with the progression of cartilage degeneration.

### 2. Expansion of BMP ligand expression drives the spread of BMP signaling in aging AC

To investigate the molecular mechanisms underlying age-associated BMP activation in mice AC, we revisited the developmental program of the secondary ossification center (SOC). Postnatally, the SOC develops through hypertrophic differentiation of epiphyseal chondrocytes and progressively advances toward the joint surface (Fig.S2A)[42,43]. This process normally terminates upon reaching adulthood, leaving a stable layer of non-hypertrophic cells, adult AC. In OA, however, as per previous data (Fig.1C-1C′′′′), chondrocyte hypertrophy expands throughout and covers the entire aging AC. We therefore sought to determine whether chondrocyte hypertrophy in OA represents a reactivation of a developmentally arrested program in adult cartilage. We analysed the expression of the BMPR1A receptor, its ligands, and the downstream BMP signaling readout pSMAD1/5/9, and assessed their relationship with the hypertrophic marker Col X across different time points in juvenile and aged AC.

Initially, we examined the presence of BMP signaling readout pSMAD1/5/9 in the AC at P15 and P30. Interestingly, immunoreactivity against pSMAD1/5/9 was detected only in the deep zone, in the vicinity of the hypertrophic chondrocytes at P15 and its expression domain got narrower as the mice reached P30 (Fig.S2B-2B′′′). To further investigate the potential reasons for the confinement of active BMP signaling to the deep zone of juvenile AC, we examined the expression patterns of BMP ligands and their receptors. Deep zone of AC at P15 showed strong immunoreactivity against the BMP2, which was further restricted by P30. Interestingly, the expression domain of BMP2 was similar to that of pSMAD1/5/9 (Fig.S2C-S2C′′′) while BMPR1A receptor expression was widespread across the AC at P15 (Fig.S2E-S2E′′′). Therefore, only the deep zone of developing AC, where both BMP ligands and receptors are present, exhibited active BMP signaling. Moreover, BMPR1A expression progressively declined in the deep zone by P30, coinciding with an increasing proportion of hypertrophic chondrocytes between P15 and P30. Moreover, Safranin O/Fast Green staining also revealed a reduction in Safranin O-stained AC thickness, accompanied by expansion of the hypertrophic region between P15 and P30 (Fig.S2F-S2F′′′). Collectively, these results indicate that in developing epiphyseal cartilage, BMP ligand exposure activates BMP signaling, driving chondrocyte hypertrophy. These hypertrophic chondrocytes subsequently show reduced BMPR1A expression.

Subsequently, we analysed the expression patterns of BMP2, BMP4 ligands and BMPR1A receptor in aging AC to assess whether BMP signaling activation in adult cartilage happens due to expansion of its ligand expression from 6 months to 3 years of age in mice. We observed minimal immunoreactivity against BMP2 and BMP4 ligands in the AC of 6-month-old mice (Fig.2B,2E and 2C, 2F, respectively). Their expression significantly increased from 18 months, had a peak between 24 and 30 months, and then declined by 36 months (Fig.2B-2B′′′′, 2E and Fig.2C-2C′′′′, 2F). On the other hand, BMPR1A immunoreactivity in the AC was highest at 6 months, steadily decreased from 18 months onward, and reached a minimum by 36 months (Fig. 2D–2D′′′′ and Fig.2G).

**Fig. 2.**
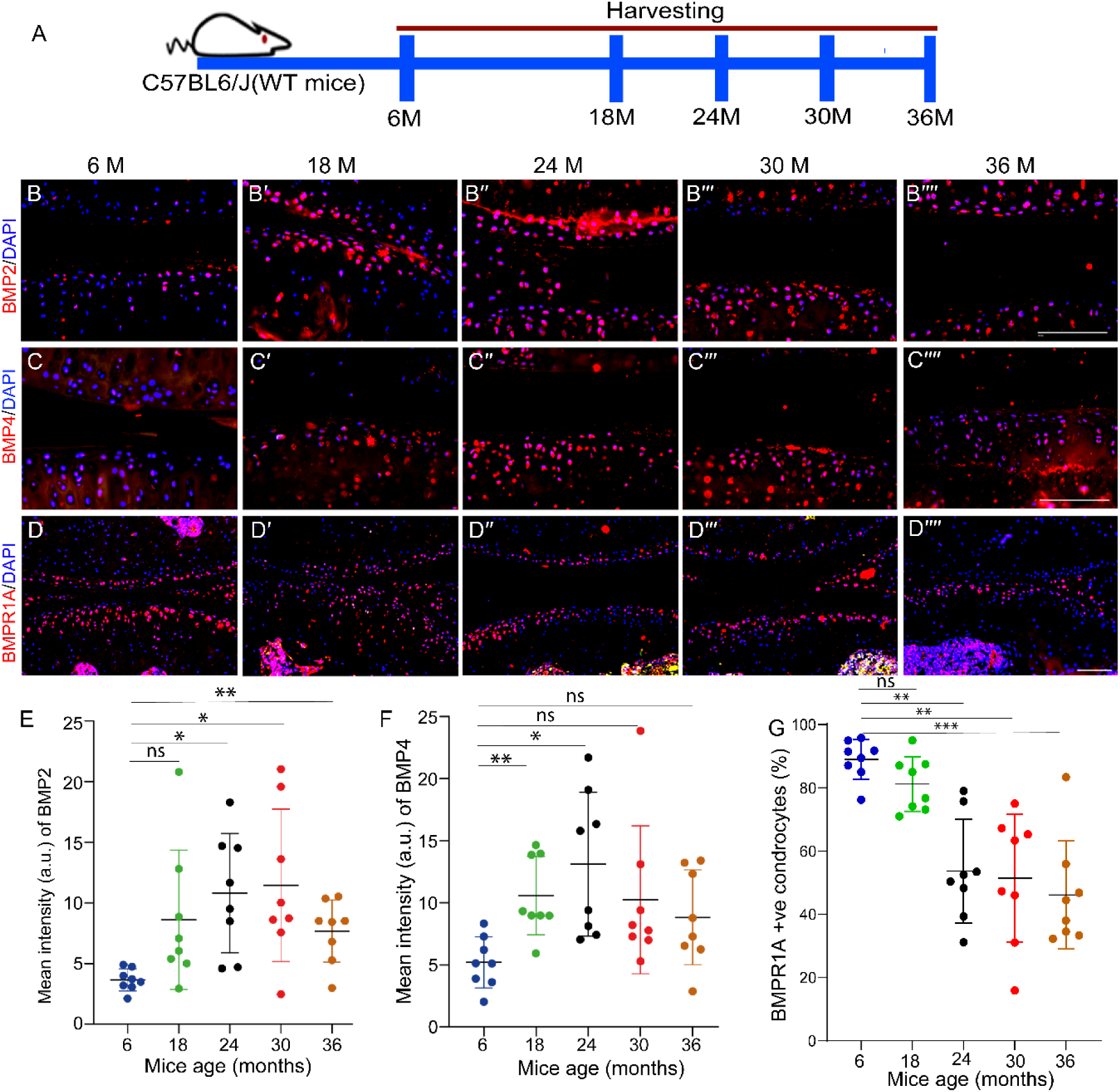
Expansion of BMP ligand expression drives the spread of BMP signaling in aging AC. (A) Schematic representation depicting selected experimental time points for analysis of BMP2, BMP4, and BMPR1A of aging C57BL/6J WT mice. (B-D′′′′) Longitudinal sections through the adult knee joints of C57BL/6J (WT mice) harvested at 6 months (B-D), 18 months (B′-D′), 24 months (B′′-D′′), 30 months (B′′′-D′′′), and 36 months (B′′′′-D′′′′) of age. Immunoreactivity for BMP2 (B-B′′′′), BMP4 (C-C′′′′), and BMPR1A (D-D′′′′). (E) Quantification data for BMP2, Brown-Forsythe, and Welch ANOVA were performed across the five sets. We compared the mean intensity of 6 months vs. 18 months; p = 0.1276 (ns), 6 months vs. 24 months; p = 0.0133 (*), 6 months vs. 30 months; p = 0.0295(*), and 6 months vs. 36 months; p =0.0081 (**). (F) Quantification data for BMP4, Brown-Forsythe, and Welch ANOVA were performed across the five sets. We compared the mean intensity of 6 months vs. 18 months; p = 0.0055 (**), 6 months vs. 24 months; p = 0.0181 (*), 6 months vs. 30 months, p = 0.1461(ns), and 6 months vs. 36 months, p =0.1126 (ns). (G) Quantification data for BMPR1A, Brown-Forsythe, and Welch ANOVA were performed across the five sets. We compared the mean intensity of 6 months vs. 18 months; p = 0.1751 (ns), 6 months vs. 24 months; p = 0.0010 (**), 6 months vs. 30 months; p = 0.0030 (**), and 6 months vs. 36 months; p =0.0003 (***). Scale bar = 100 µm; n=8 per group. Error bars represent the mean ± standard deviation (S.D.).

Together, these data indicate that articular chondrocytes are intrinsically competent to execute BMP signaling even at a young age (6 months) but fail to do so due to limited BMP ligand availability. With aging, expanded BMP ligand expression activates BMP signaling, driving chondrocyte hypertrophy, subsequent BMPR1A depletion, and the progression of OA. Notably, this hypertrophic response in aging AC reflects reactivation of developmental molecular programs that are normally silenced in healthy adult mice.

### 3. Local intra-articular delivery of BMP ligands is sufficient to induce cellular and molecular events of age-induced OA

Our previous results suggested that the availability of BMP ligands are the primary limiting factor driving chondrocyte hypertrophy and, consequently, age-associated OA (Fig.2). Therefore, to confirm it further, we administered two doses of recombinant human BMP2 protein/vehicle (500 ng/dose) into the joint cavities of 6-month-old mice and collected the tissue 48 hours after the second injection (Fig.S3A). A drastic up-regulation of nuclear pSMAD1/5/9 in the BMP2-injected samples as compared to the vehicle-injected samples suggests that the local delivery of BMP ligand is sufficient to activate BMP signaling in the healthy mice AC (Fig.S3B-3B′′). In parallel, another group of mice treated with BMP2/vehicle were sacrificed on post-injection days 28 and 56 (Fig.3A). We examined the OA-like changes in the AC samples obtained from mice injected with BMP2 protein. At 28- and 56-day post-injection, immunohistochemistry results revealed that samples injected with BMP2 had significantly depleted native Col II compared to those injected with the vehicle (Fig.3B-3B′′,3H). Further, in the BMP2-injected samples, there was a significant increase in the number of Col X-expressing hypertrophic chondrocytes compared to the samples that had been treated with the vehicle (Fig.3C-3C′′,3I). Earlier results indicated that hypertrophic chondrocytes deplete BMPR1A receptor expression during juvenile AC development (Fig. S2E-S2E′′′) or age-induced OA (Fig. 2D-2D′′′′). Immunohistochemistry against BMPR1A in the BMP2-injected sample revealed a substantial decrease in the number of BMPR1A-expressing chondrocytes compared to the vehicle-treated samples (Fig.3D-3D′′,3J). Further, dual immunohistochemistry against Col X and BMPR1A in BMP2/Vehicle-treated samples clearly showed that cells that express Col X are devoid of BMPR1A and vice versa (Fig.3E-3E′′) except for a few that exhibit minimal immunoreactivity for both markers. The presence of proteoglycans is a key measure for assessing the integrity of AC, which was depleted in OA cartilage. We analysed the status of proteoglycan by Safranin O /Fast green staining of samples injected with BMP2 protein and Vehicle. Strong safranin O staining in the vehicle-injected samples revealed preserved AC integrity, while diminished safranin O in BMP2-injected samples showed loss of AC integrity (Fig.3F-3F′′,3G-3G′′). Thus, intra-articular BMP2 delivery is sufficient to induce BMP signaling in 6-month-old mice, driving the onset of cellular and molecular events of age-associated OA, irrespective of the mice’s chronological age.

**Fig. 3.**
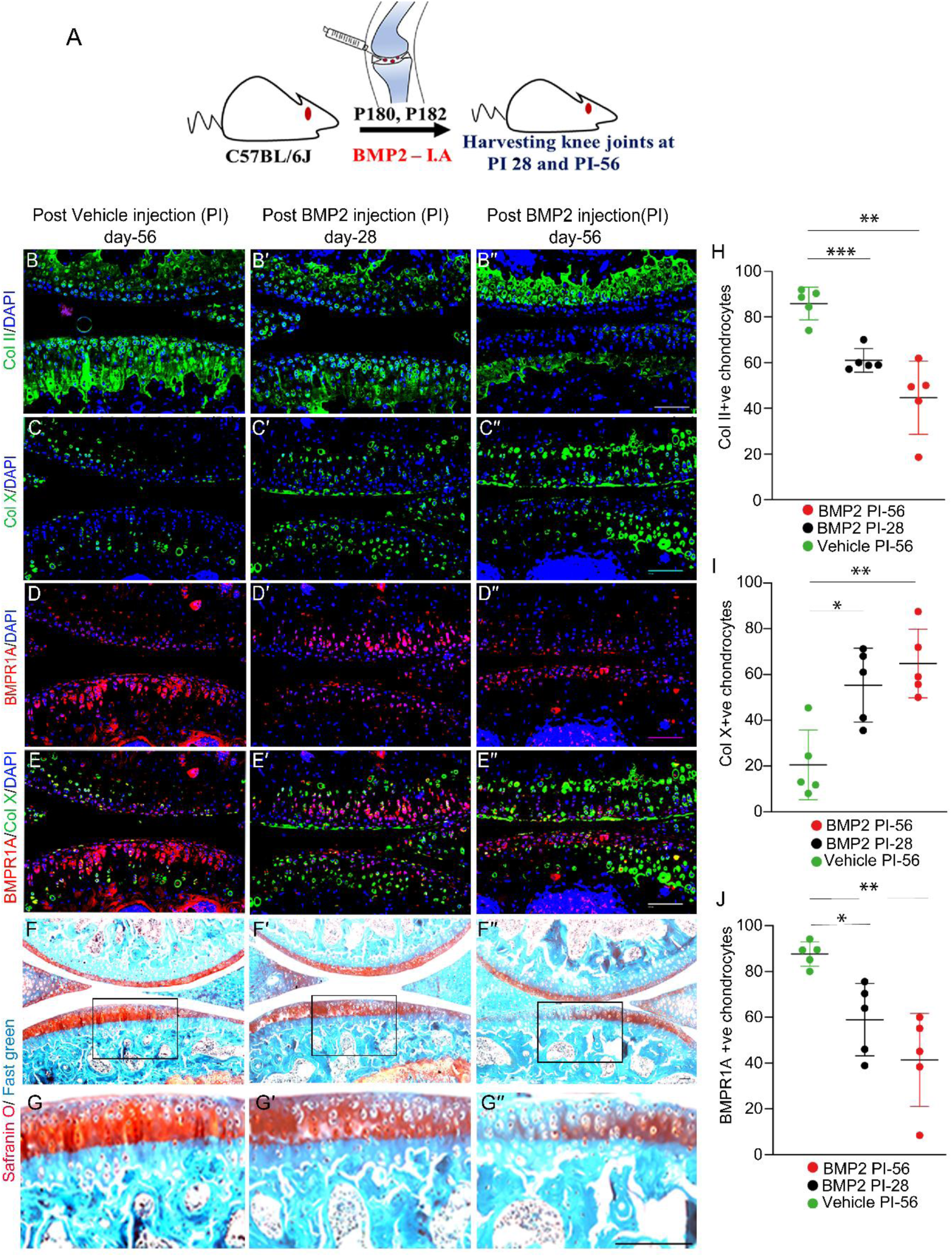
Local intra-articular delivery of BMP ligands is sufficient to trigger the cellular and molecular events of age-associated OA in young mice. (A) Schematic for intra-articular injection of recombinant BMP2 protein in C57BL/6J (WT) mice at 6 months of age. (B-G′′) Longitudinal sections through the adult knee joints of C57BL/6J (WT mice) harvested post-vehicle injection (PI Day-56) (B-G), 28 days post-BMP2 injection (PI Day-28) (B′-G′), 56 days post-BMP2 injection (PI Day-56) (B′′-G′′). Immunoreactivity for ColII (B-B′′), Col X (C-C′′), BMPR1A (D-D′′), dual immunostaining for BMPR1A (red) and Col X (green) (E-E′′), Safranin O staining (F-F′′) and its inset (G-G′′, Scale bar:20µm); (H) Quantification data for ColII, with Brown-Forsythe and Welch ANOVA was performed along the three sets. We compared the mean of vehicle PI-56 vs. BMP2 PI-28; p = 0.0007 (***), vehicle PI-56 vs. BMP2 PI-56; p = 0.0044 (**). (I) Quantification data for Col X, Brown-Forsythe, and Welch ANOVA were performed on the three sets. We compared the mean of vehicle PI-56 vs. BMP2 PI-28; p = 0.0144 (*), vehicle PI-56 vs. BMP2 PI-56; p = 0.0031 (**); (J) Quantification data for BMPR1A, Brown-Forsythe, and Welch ANOVA were performed along the three sets. We compared the mean of Vehicle PI-56 vs. BMP2 PI-28; p = 0.0219 (*), Vehicle PI-56 vs. BMP2 PI-56; p = 0.0098 (**). Scale bar = 100 µm; n=5 per group. Error bars represent the mean ± standard deviation (S.D.).

### 4. Aging AC exhibits reduced β-catenin and elevated ICAT, a downstream target of BMP signaling

The Wnt signaling pathway plays a crucial role in the differentiation of AC during the embryonic development [27,28]. However, both gain and loss of function mutation of the canonical Wnt pathway resulted in the deterioration of adult mouse AC integrity[30–32]. Therefore, we were curious to examine the exact role of Wnt-β-catenin signaling in the maintenance of AC during aging. We harvested knee joints from mice aged 6 to 36 months and performed immunohistochemistry to examine the status of canonical Wnt signaling readout β-catenin (Fig.4A). Strong immunoreactivity against nuclear β-catenin was detected in 6-month-old mice, which decreases with age starting at the 18th month and approaches a minimum in 36-month-old AC (Fig.4B-B′′′′ and 4I).

**Fig. 4.**
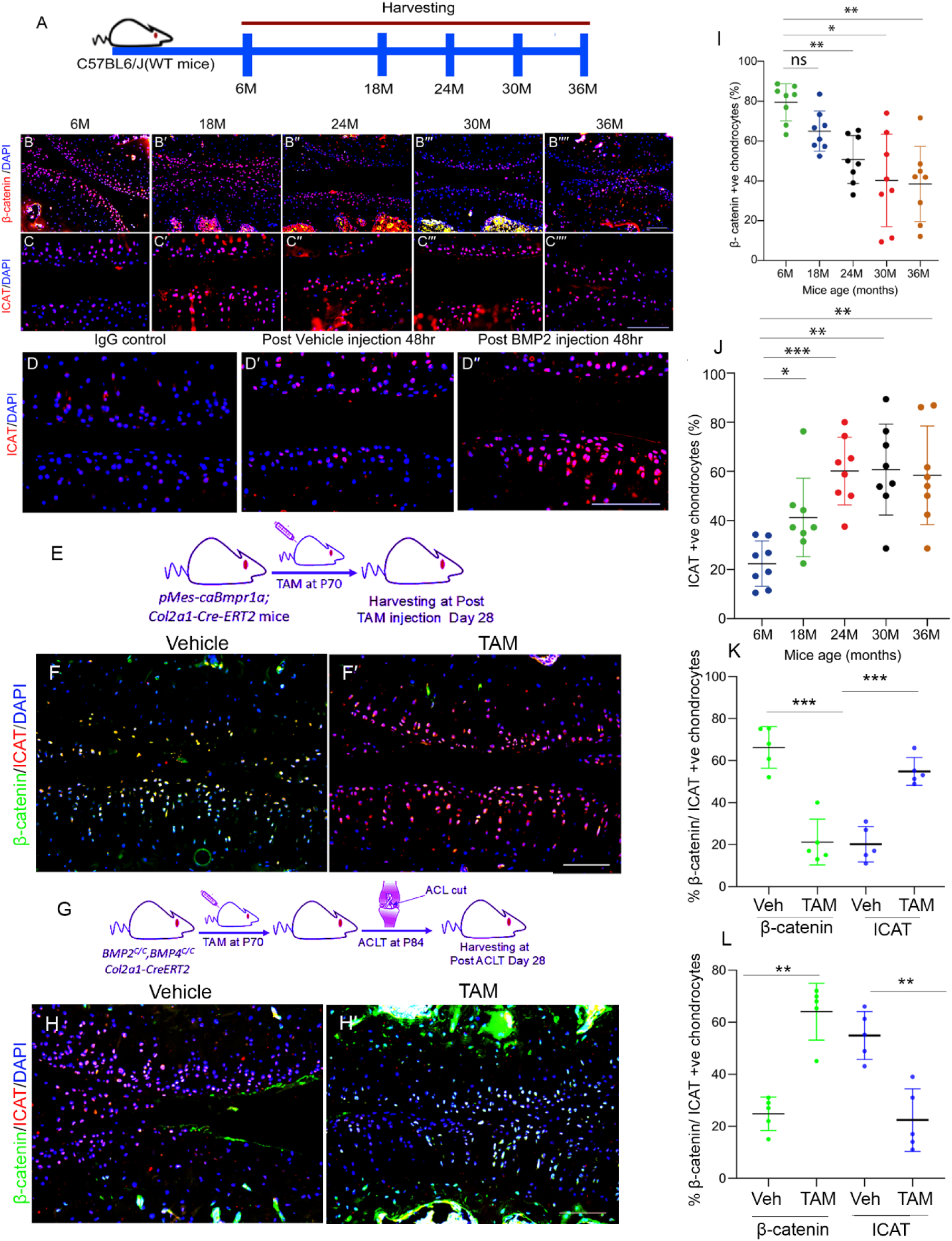
Aging AC exhibits reduced β-catenin and elevated ICAT, a downstream target of BMP signaling. (A) Schematic representation depicting selected experimental time points for analysis of β-catenin and ICAT in aging C57BL/6J WT mice. (B-C′′′′) Longitudinal sections through the adult knee joints of C57BL/6J (WT mice) harvested at 6 months (B-C), 18 months (B′-C′), 24 months(B′′-C′′), 30 months (B′′′-C′′′), and 36 months (B′′′-C′′′) of age. Immunoreactivity for β-catenin (B-B ′′′′), ICAT (C-C′′′′). (D-D′′) Longitudinal sections through the adult knee joints of C57BL/6J (WT mice) were harvested post-vehicle injection 48 hours (D′), post-BMP2 injection 48 hours (D′′), and IgG control (D). (E) Experimental design for BMP gain-of-function using pMes-caBmpr1a; TgCol2a1-Cre-ERT2 mice. Immunohistochemistry for β-catenin and ICAT was performed at 28 days post-TAM injection in Vehicle (F) and TAM-treated (F′) groups. (G) Experimental strategy for BMP loss-of-function using Bmp2^c/c^; Bmp4^c/c^; TgCol2a1-Cre-ERT2 mice. Immunohistochemistry for β-catenin and ICAT was conducted 28 days after TAM injection in Vehicle (H) and TAM-treated (H′) groups. (I) Quantification data for β-catenin, Brown-Forsythe, and Welch ANOVA were performed across the five sets. We compared the means of 6 months vs. 18 months; p = 0.0640 (ns), 6 months vs. 24 months; p = 0.0010 (**), 6 months vs. 30 months, p = 0.0101(*) and 6 months vs. 36 months, p =0.0017(**); (J) Quantification data for ICAT, Brown-Forsythe and Welch ANOVA were performed along the five sets. We compared the means of 6 months vs. 18 months; p = 0.0459 (*), 6 months vs. 24 months; p = 0.0001 (***), 6 months vs. 30 months p = 0.0012(**) and 6 months vs. 36 months p =0.0034(**). (K) Quantification of β-catenin and ICAT in BMP gain-of-function samples using *pMes-caBmpr1a; TgCol2a1-Cre-ERT2* mice following TAM or Vehicle injection. Brown–Forsythe and Welch ANOVA were used for statistical analysis. β-catenin: Vehicle vs TAM, p = 0.0006(***); ICAT: Vehicle vs TAM, p =0.0005(***). (L) Quantification of β-catenin and ICAT in BMP loss-of-function samples using Bmp2^c/c^; Bmp4^c/c^; TgCol2a1-Cre-ERT2 mice injected with TAM or Vehicle injection following ACLT surgery. Brown–Forsythe and Welch ANOVA were used for statistical analysis. β-catenin: Vehicle vs TAM, p = 0.0013(**); ICAT: Vehicle vs TAM, p = 0.0070(**). Scale bar = 100 µm. *n* = 8 per group for graphs I and J; *n* = 5 per group for graphs K and L. Error bars represent mean ± standard deviation (S.D.).

Given the age-associated increase in BMP and decrease in Wnt/β-catenin activity (Fig.1F-F′′′′ and Fig.4B-B′′′′), we investigated potential regulators underlying their inverse expression profiles and assessed whether these pathways are functionally interconnected or operate independently. Moreover, literature suggests that BMP signaling regulates the Wnt pathway in colorectal cancer (CRC) tissue via modulating its downstream protein ICAT (inhibitor of β-catenin and TCF-4), a known canonical Wnt signaling inhibitor[44]. Therefore, we performed immunohistochemistry against ICAT on AC samples from mice aged 6 months to 36 months. Minimal immunoreactivity against ICAT was observed in 6- month-old mice, which progressively increased after the 18th month and reached its highest at 36 months (Fig.4C-C′′′′,4J).

This evidence implies that as mice age, an increased BMP signaling possibly triggers the production of ICAT, which subsequently inhibits Wnt/β-catenin signaling in AC. However, it was unclear whether ICAT expression is directly regulated by BMP signaling or has increased just because of aging. Therefore, BMP signaling was activated in AC of 6-month-old mice by injecting BMP2 protein/Vehicle directly into the joint cavity. Tissues were collected 48h post BMP2/vehicle injection to evaluate ICAT status in AC. Strong immunoreactivity against ICAT was found in BMP-2-injected samples, while it was minimally detected in the vehicle control group (Fig.4D-D′′). This data suggests that BMP signaling directly regulates ICAT expression, which suppresses the Wnt-β-catenin pathway during age-associated OA. To further validate this, we genetically activated BMP signaling in postnatal cartilage using tamoxifen (TAM)-induced *pMes-caBmpr1a; TgCol2a1-Cre-ERT2 mice*. TAM or vehicle was administered at P70, and tissues were collected 28 days after injection to evaluate ICAT and β-catenin expression (Fig.4E). A markedly increased ICAT immunoreactivity was observed in the TAM-injected group, whereas only minimal staining was detected in the vehicle-treated samples (Fig.4F-4F′′ and Fig.4K). In contrast, β-catenin expression was reduced in the TAM-injected group compared to the vehicle-treated group, demonstrating an inverse relationship between ICAT and β-catenin expression in these samples (Fig.4F-4F′ and Fig.4K).

Previous findings suggest that elevated BMP signaling plays a regulatory role in the development of PTOA[24]. To determine whether BMP signaling is required for ICAT expression, we utilised *Bmp2 ^c/c^; Bmp4 ^c/c^; TgCol2a1-Cre-ERT2* mice. In this model, BMP2 and BMP4 are conditionally ablated in a tissue-specific manner upon intraperitoneal TAM administration. TAM or vehicle was injected at P70, followed by ACLT surgery at P84 to trigger BMP signaling-induced OA. Expression of ICAT and β-catenin was analysed in these samples at post ACLT day 28. Strong ICAT immunoreactivity was observed in the vehicle-treated group, while only minimal staining was detected in the TAM-treated BMP-depleted group (Fig.4H-4H′ and Fig.4L). In contrast, β-catenin showed stronger immunoreactivity in the TAM-injected samples compared to the vehicle-treated group (Fig.4H-4H′ and Fig.4L). Collectively, these results strongly suggest that BMP-induced ICAT expression directly inhibits canonical Wnt/β-catenin signaling in aging AC during OA progression.

### 5. Constitutive Wnt/β-catenin signaling fails to protect against OA due to consequential activation of BMP signaling

Previous results suggest that the level of Wnt/β-catenin signaling in the AC decreases with aging of the AC (Fig.1F-F′′′′ and Fig.4B-B′′′′). Thus, a logical extension would have been that activating Wnt/β-catenin signaling can mitigate OA. However, the existing literature demonstrates that promoting Wnt/β-catenin signaling induces OA-like changes in the adult AC[30,32]. To resolve this apparent contradiction, we delved deeper to know the precise role of Wnt/β-catenin in maintenance of adult AC.

To this end, we employed the *Catnb^lox(ex3)/ (ex3)^*; *TgCol2a1: Cre-ERT2* mouse strain, in which Wnt/β-catenin could be activated upon intra-peritoneal injection of TAM (Fig.5A). We analysed the effect of constitutively activated Wnt/β-catenin signaling in adult AC. TAM or vehicle was administered at P70, and tissues were collected 28 days post-injection for molecular analyses. Notably, compared with vehicle-treated controls, TAM-induced activation of Wnt/β-catenin signaling led to decreased expression of the AC marker Col II and increased expression of the hypertrophic marker Col X (Fig. S4). These unexpected results suggest that constitutive activation of Wnt/β-catenin signaling resulted in cartilage degeneration rather than preservation, despite the established link between reduced Wnt/β-catenin activity and age-related OA progression. To explore the potential mechanisms underlying this effect, we next analysed the status of BMP signaling in these samples. Interestingly, as early as 14 days following TAM injection, pSMAD1/5/9 immunoreactivity was markedly increased in the TAM-induced group compared with vehicle-treated groups. Moreover, elevated pSMAD1/5/9 levels persisted in the AC up to 35 days after Wnt/β-catenin activation (Fig. 5B-C′′). Given the OA-like features of these tissues and the known association of OA with inflammation, we also analysed the expression of inflammation-related genes[45]. We found that nuclear NF-κB levels were significantly elevated in the TAM-injected group compared with vehicle-injected controls (Fig.5D-5D′, 5E-5E′ and Fig.5F). To further investigate whether NF-κB activation arises directly from Wnt/β-catenin signaling or occurs indirectly through BMP signaling. We cultured BRITER cells[46] under chondrogenic conditions and exposed them to recombinant BMP2 protein or vehicle for 48h, and immunocytochemistry was performed against NF-κB. An elevated immunoreactivity for NF-κB was observed in BMP2-treated cells compared to controls, indicating that elevated BMP signaling can directly regulate nuclear NF-κB activity in adult AC (Fig.6H-H′′). Taken together, these results suggest that although Wnt/β-catenin signaling is essential for maintaining AC, its constitutive activation promotes cartilage degeneration through BMP signaling. Consequently, targeting BMP signaling, rather than activating Wnt/β-catenin, may represent an effective pharmacological strategy for managing age-associated OA.

**Fig. 5.**
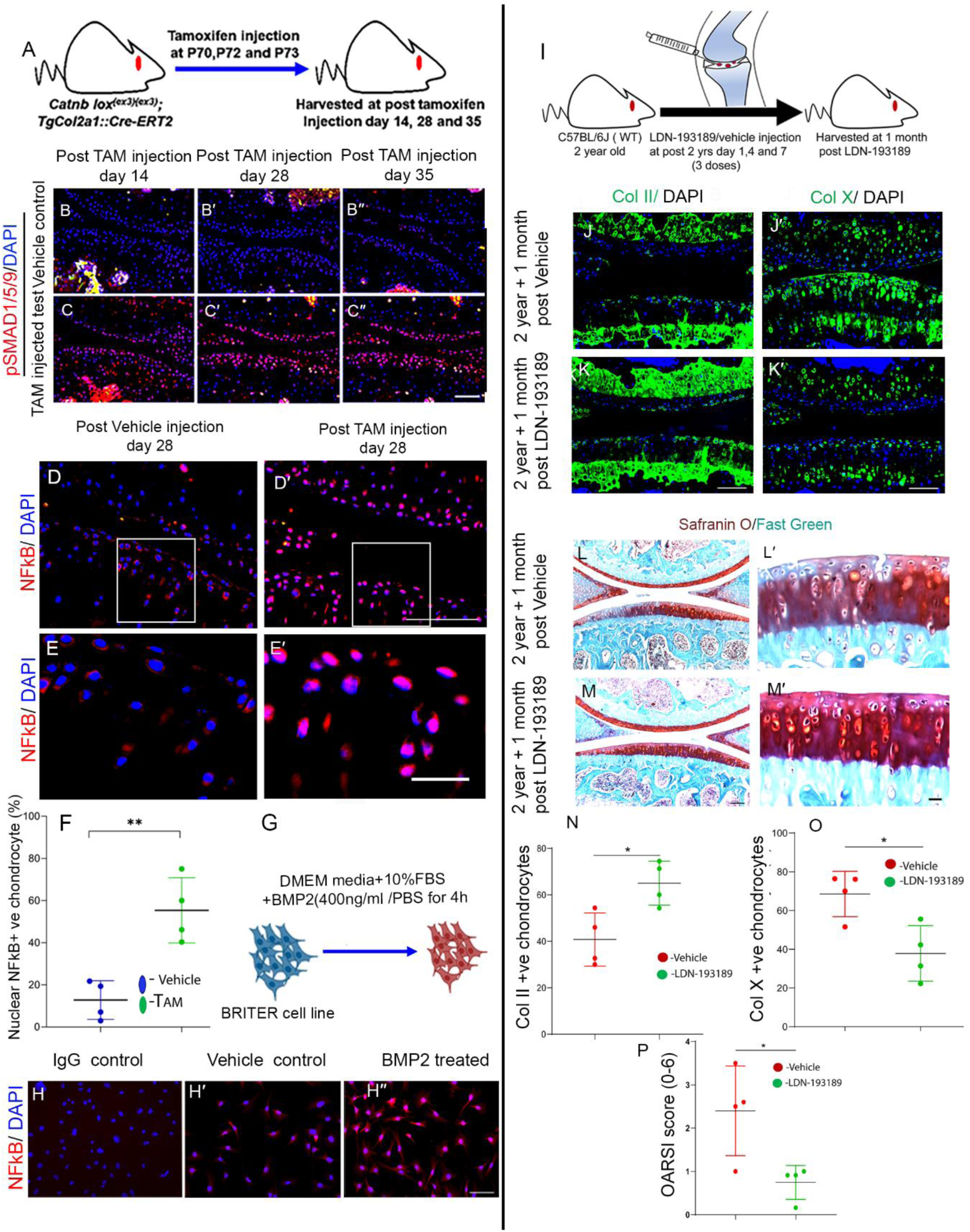
Constitutively active Wnt-β-catenin deteriorates AC by enhancing active BMP signaling. (A) Schematic representation of activation of Wnt-β-catenin in *Catnb^lox(ex3)/lox(ex3)^; TgCol2a1: Cre-ERT2* mice via intra-peritoneal tamoxifen (TAM) injection at P70. (B-C′′) longitudinal section of the knee joint of *Catnb^lox(ex3)/lox(ex3)^; TgCol2a1: Cre-ERT2* mice post-vehicle/tamoxifen (TAM) injection Day 14 (B, C), Day 28 (B′ C′) and Day 35 (B′′, C′′). Immunohistochemistry against pSMAD1/5/9 vehicle-injected (B, B′ and B′′) and TAM-injected group (C, C′ and C′′). (D-E′) Immunohistochemistry against NFκB in vehicle-injected (D and E), and TAM-injected (D′ and E′). E and E′ are the insets of D and D′, respectively. (F) Quantification data for Nuclear NFκB in articular chondrocyte. The Unpaired t-test with Welch’s correction was performed on the two sets. We compared the means of Vehicle vs TAM injected sample; p=0.0055 (**); (G) Schematic representation for treatment of BRITER cell line with BMP2 protein and immunoreactivity for NFκB (H-H′′); of IgG control (H), 4 hours post-vehicle treatment (H′) and 4 hours post-BMP2 treatment (H′′). (I) Schematic representation of regimen for local inhibition of BMP signaling using LDN-193189 of C57BL/6J (WT mice); (J-M) Longitudinal sections through the adult knee joints of C57BL/6J (WT mice) harvested at 2 years and 1-month post-vehicle injection (J-J′ and L-L′) and 1-month post-intra-articular injection of LDN-193189 (K-K′ and M-M′); Immunoreactivity for ColII (J and K), Col X (J′-K′); Safranin O staining (L-M), inset (L′-M′ inset scale bar: 20 µm) (N) Quantification data for ColII, Unpaired t-test with Welch’s correction was performed along the two sets. We compared the means of Vehicle vs. LDN-193189; p=0.0172 (*); (O) Quantification data for Col X, Unpaired t-test with Welch’s correction was performed along the two sets. We compared the means of Vehicle vs. LDN-193189; p= 0.0180 (*); (P) OARSI score, Unpaired t-test with Welch’s correction was performed along the two sets. We compared the means of Vehicle vs. LDN-193189; p=0.0425 (*). Scale bar = 100 µm; n=4 per group. Error bars represent the mean ± standard deviation (S.D.).

**Fig. 6.**
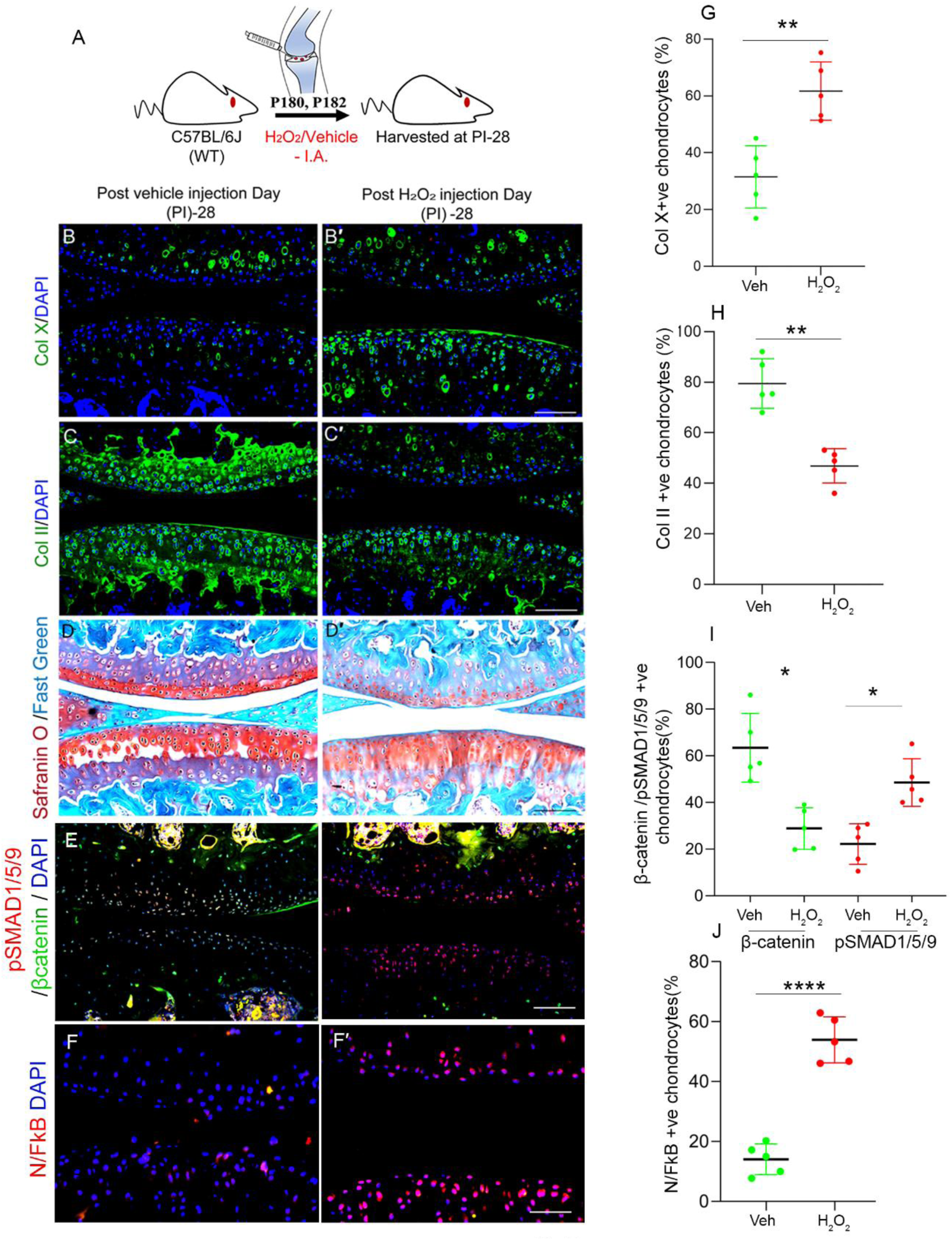
Local supply of H_2_O_2_ in the joint cavity of mice induces OA-like phenotype. (A) Schematic for intra-articular injection of H_2_O_2_/ Vehicle control at P180, P181(2 doses), and harvested post-injection 28 days. (B-D′) Longitudinal sections through the adult knee of 28 days post-vehicle injection (B-D), 28 days post-H_2_O_2_ injection (B′-D′); Immunoreactivity for Col X (B-B′), ColII (C-C′), Safranin O staining (D-D′), Psmad1/5/9 and β-catenin(E-E′), NFκB (F-F′). (G) Quantification of Col X. An unpaired *t*-test with Welch’s correction was performed to compare Vehicle and H₂O₂-treated groups; p= 0.0013(**) (H) Quantification of Col II. An unpaired *t*-test with Welch’s correction was performed to compare Vehicle and H₂O₂-treated groups; p=0.0011(**) (**I)** Quantification of pSMAD1/5/9 and β-catenin. Brown-Forsythe and Welch ANOVA were used to compare Vehicle and H₂O₂-treated groups; β-catenin, p=0.0133(*) and pSMAD1/5/9 p=0.0106 (*) (J). Quantification of NF-κB. An unpaired *t*-test with Welch’s correction was performed to compare Vehicle and H₂O₂-treated groups; p<0.0001(****). Error bars represent the mean ± standard deviation (S.D.). Scale bar = 100 µm, n = 5 per group.

To test our hypothesis, we administered 450 picograms of the BMP signaling inhibitor LDN-193189 into the joint cavity of 2-year-old C57BL/6J mice, with the contralateral limb receiving vehicle injections. Three doses of LDN-193189 were given at three-day intervals, and tissues were collected one month later (Fig. 5I). Analysis of AC revealed an increase in Col II and a decrease in the hypertrophic marker Col X expression in the LDN-193189-treated group compared to group treated with vehicle only (Fig. 5J-5K, 5N; 5J′–5K′, 5O). Furthermore, Safranin O/Fast Green staining demonstrated improved integrity with fewer vertical clefts in the LDN-193189-treated group compared with the vehicle-treated group (Fig. 6L–6M, 6L′–6M′). Consistently, OARSI scoring revealed lower grades in the LDN-193189–injected joints, indicating effective protection against age-associated OA (Fig. 6P).

### 6. ROS acts as an upstream driver of BMP signaling to induce OA

Next, we aimed to explore potential triggers behind elevated BMP ligand expression, thereby activating BMP signaling in aging AC. The mouse embryos were pulse-labelled with BrdU at E12.5, allowed to develop to birth, and the offspring were maintained for 2 and 6 months before knee joint tissues were collected and subjected to immunohistochemistry against BrdU (Fig. S5). Surprisingly, BrdU-labelled cells were detected throughout the AC, indicating that AC cells possess a remarkably long lifespan. With advancing age, chondrocytes experience increased oxidative stress due to mitochondrial dysfunction, reduced antioxidant capacity, and cumulative mechanical and metabolic stress[47,48]. Moreover, increased ROS production is closely associated with enhanced BMP signaling in articular chondrocytes, with evidence suggesting a reciprocal regulatory relationship between ROS and BMP signaling[49,50]. To test whether ROS activates BMP signaling in aging AC, 2 alternate doses of 2% H₂O₂(v/v) or vehicle were injected into the knee joints of 6-month-old mice (Fig. 6A). Tissues were collected 28 days later, and OA-like changes were analysed in these samples. Strong immunoreactivity for Col X was observed in the H₂O₂-treated samples, whereas only minimal staining was detected in the vehicle-treated controls (Fig.6B-B′ and Fig.6G). The increased Col X expression in the H₂O₂-treated samples clearly indicates chondrocyte hypertrophy, a characteristic hallmark of OA cartilage. Moreover, immunoreactivity for the native cartilage marker Col II was reduced in the H₂O₂-treated samples, whereas it was well maintained in the vehicle-treated group (Fig.6C-C′ and Fig.6H). The increased proportion of Col X-expressing hypertrophic chondrocytes, along with the decrease in Col II expression, clearly suggests a shift in the fate of articular chondrocytes toward OA progression. To evaluate overall cartilage integrity, we performed Safranin O/Fast Green staining of these samples. The H₂O₂-treated samples showed a marked reduction in Safranin O staining intensity and rough surface compared with the vehicle-treated controls, indicating significant loss of proteoglycans, thereby compromising cartilage integrity (Fig.6D-D′and Fig.6H). These fundamental alterations in AC following H₂O₂ treatment strongly suggest that reactive ROS are sufficient to trigger OA-like changes.

Previous results revealed that OA-like changes are induced by increased BMP and decreased Wnt/β-catenin signaling (Fig.1F-F′′′′ and Fig.4B-B′′′′). To determine whether the ROS-induced OA-like changes are driven by activation of BMP signaling, which could suppress Wnt/β-catenin signaling through ICAT. We performed dual immunohistochemistry for the BMP signaling readout pSMAD1/5/9 and β-catenin in these samples. Notably, H₂O₂-treated samples showed increased pSMAD1/5/9 and decreased β-catenin immunoreactivity compared to the vehicle-treated group (Fig.6E-E′ and Fig. 6I). Additionally, ICAT expression was elevated in the H₂O₂-treated group (Fig.S7). Moreover, earlier results also revealed that increased BMP signaling can also induce nuclear NF-κB signaling (Fig.5H-H′′). Therefore, we examined the status of NF-κB in these samples. Strong immunoreactivity for NF-κB was observed in the H₂O₂-treated samples, whereas it was only minimally detected in the vehicle-treated controls (Fig.6F-F′ and Fig. 6J).

These results strongly suggest that elevated ROS in aging chondrocytes serves as a key driver of BMP signaling, leading to AC degeneration by inhibiting Wnt/β-catenin signaling via ICAT. Furthermore, increased BMP signaling promotes inflammation through nuclear translocation of NF-κB. Thus, ROS may act as a primary trigger for the cascade of events driving degeneration in aging AC.

## Discussion

Old age is the primary risk factor for OA[16,47]. To understand its molecular etiology, we used a suitable model system to investigate its underlying mechanism through detailed analyses of temporal histological, molecular, and radiological changes in aging AC. We observed that during early developmental stages, BMPR1A receptors were uniformly expressed throughout the developing epiphyseal cartilage, whereas BMP ligands were initially confined to the center. As development proceeded, the BMP ligand expression domain expanded toward the outer layers, inducing hypertrophy in BMP-exposed chondrocytes. This pattern persisted throughout development and contributed to the formation of the SOC center. Subsequently, upon reaching adulthood, termination of BMP ligand expression maintains adult AC cells in a non-hypertrophic state. However, during age-associated OA, the normally restricted BMP ligand expression expands across the AC, activating BMP signaling and driving chondrocyte hypertrophy in the entire tissue. Moreover, local delivery of BMP2 protein into the joint cavity of 6-month-old healthy AC was found to be sufficient to trigger cellular and molecular events of aging cartilage independent of chronological age. Furthermore, chondrocytes lose BMPR1A expression upon hypertrophic differentiation, a pattern that is observed both during endochondral ossification and in aging AC. This absence of BMPR1A in hypertrophic chondrocytes appears to be an intrinsic property of these cells. This likely explains the biphasic pattern seen in aging AC, in which the proportion of chondrocytes exhibiting active BMP signaling initially increases and then declines. A similar mechanism may underlie the peak and subsequent reduction of BMP signaling observed during endochondral bone formation as chondrocytes differentiate to hypertrophic[20,51]; however, BMPR1A expression in this context has yet to be formally evaluated.

Wnt/β-catenin signaling is a well-established pathway that regulates AC differentiation during embryonic development [27,28]. Our data further suggest that AC cells exhibit active Wnt/β-catenin signaling, which progressively declines with age. This decline may explain why Wnt loss of function promotes OA in young adults [32]. However, gain-of-function activation of Wnt/β-catenin signaling not only fails to protect against OA but actively promotes OA-like changes. Further analysis of these tissues revealed that constitutive activation of Wnt/β-catenin signaling induces BMP signaling in AC, thereby promoting chondrocyte hypertrophy. [30,31]. These findings suggest that while basal Wnt/β-catenin activity is indispensable for AC homeostasis, since its loss results in an OA-like phenotype, its sustained hyperactivation paradoxically triggers BMP signaling, thereby driving cartilage degeneration rather than conferring protection. Gil et *al.* also emphasise that maintaining optimal Wnt/β-catenin signaling is essential, and any deviation from it can have detrimental effects on cartilage integrity *(54).* However, maintaining an optimal level of Wnt/β-catenin signaling at the cellular level is challenging through pharmacological or genetic approaches, as its activity in degenerating AC is not simply binary but instead varies among individual cells within the tissue. Hence, inhibition of BMP signaling rather than modulation of Wnt/β-catenin signaling seems a practical approach to develop disease-modifying therapy for OA.

Furthermore, while young healthy cartilage exhibits active Wnt/β-catenin signaling with minimal BMP activity, this balance progressively shifts with aging toward enhanced BMP signaling. We identify ICAT, a BMP signaling-induced inhibitor of the Wnt/β-catenin pathway, as a key mediator of this reciprocal regulation[44]. Consistently, intra-articular BMP administration or genetic activation of BMP signaling increased ICAT expression, whereas deletion of BMP2 and BMP4 in adult cartilage abolished ICAT induction in the PTOA. Overall, these findings support the existence of a BMP-dependent negative feedback mechanism that modulates Wnt/β-catenin signaling, positioning ICAT as a pivotal mediator of BMP-Wnt crosstalk during cartilage aging and osteoarthritis progression. Nevertheless, additional genetic and biochemical studies are required to definitively confirm this regulatory pathway.

Notably, the presence of embryonically labelled BrdU-positive cells in adult AC indicates that these chondrocytes are long-lived, providing a potential cellular basis for the age-related decline in mitochondrial function, resulting in a metabolic shift from oxidative phosphorylation to aerobic glycolysis [52–54]. Further, inefficient mitochondrial activity may lead to increased ROS production in these cells[55]. Higher ROS accumulation in AC cells over time is known to trigger BMP signaling, or vice versa [50,56]. Consistently, our data suggest that H₂O₂ injection into the joint cavity of young mice elevates BMP signaling and recapitulates the cellular and molecular hallmarks of age-associated OA, independent of chronological age. These findings indicate that oxidative stress may be a key driver of the age-associated increased BMP signaling and consequent OA development. Moreover, an elevated BMP signaling also upregulates GLUT1 expression, which is associated with increased chondrocyte hypertrophy[57]. Thus, increased ROS production in aging articular chondrocytes possibly boosts BMP signaling which in turn stimulates GLUT1 expression and chondrocyte hypertrophy during age-associated OA. However, further research is needed to test these hypotheses and identify additional BMP upstream factors other than ROS in this context.

A limitation of this study is that the sex of the animals was not specifically accounted for in the age-related experiments. Although the molecular signatures we observed were largely consistent between male and female mice, subtle sex-specific differences in age-associated osteoarthritis may still exist and potentially affect disease progression or severity. However, our data indicate that the expression dynamics of OA markers were similar in both sexes.

Nonetheless, we propose that age-associated OA may represent an almost inevitable outcome, largely dictated by the kinetics of BMP ligand expression across aging AC, which varies among individuals. Elevated ROS in aging chondrocytes may act as a trigger for increased BMP ligand expression and could be further influenced by factors such as genetic background and lifestyle. Despite this inter-individual heterogeneity, targeted modulation of BMP signaling in situ represents a promising therapeutic strategy, regardless of the age at disease onset.

## Material and Method

### Study design

#### Sample size

We applied statistical power calculations using G*Power3.1 software to determine the sample size. The number of animals for every experiment was calculated based on the “Resource Equation Approach” explained by Arifin et.al. (2017[58]. Each experimental group consists of 4-8 animals, depending on the effect size of the phenotype of the experiment. For each experiment, a single animal is considered as one experimental unit.

#### Data exclusion

No data points were excluded as outliers. Animals that died before the experimental endpoint were excluded from analysis. All measurements were obtained at the predefined endpoints of each experiment.

#### Experimental design

Experiments were designed as controlled laboratory studies incorporating both in vitro and in vivo approaches. In vivo experiments utilised mouse models to examine age-associated changes in AC and the roles of BMP and Wnt/β-catenin signaling during OA progression. Experimental manipulations included intra-articular injection of BMP2 protein, H₂O₂ and LDN-193189 as well as genetic manipulations where indicated. Structural and cellular changes in joint tissues were evaluated primarily through imaging-based analyses, including μCT to assess joint structure and mineralised tissues, and microscopic imaging of histological and immunohistochemically stained sections to examine AC morphology, chondrocyte hypertrophy, and signaling activity. These imaging datasets were used for both qualitative and quantitative analyses to evaluate changes associated with aging and alterations in BMP and Wnt/β-catenin signaling.

#### Randomisation

Age- and strain-matched mice were used, and animals were randomly assigned to experimental groups where appropriate. Both sexes were included to minimise sex bias.

### Animal study protocols

All animals were bred, housed, and maintained in the Central Experimental Animal Facility (CEAF) at the Indian Institute of Technology, Kanpur, India, with food and water provided *ad libitum* under a 12-h light/12-h dark cycle. All animal experiments were conducted in accordance with protocols approved by the Institutional Animal Ethics Committee (IAEC) of IIT Kanpur (protocol numbers IITK/IAEC/2022/1167 and IITK/IAEC/2023/1174), under the guidelines of the Committee for Control and Supervision of Experiments on Animals (CPCSEA), Government of India. All experiments were performed in compliance with the ARRIVE guidelines[59].

### Generation of mouse lines

C57BL/6J wild-type mice (JAX stock no. 000664) and *TgCol2a1-CreERT2* mice (JAX stock no. 006774) were obtained from The Jackson Laboratory (USA). *Catnb^lox(ex3)/lox(ex3)^* mouse line was originally generated by Makoto Mark Taketo (Kyoto University, Japan) and was obtained through Shubha Tole (TIFR, Mumbai, India) (57). *Tg(Col2a1-CreERT2)* mice were crossed with *Catnb^lox(ex3)/lox(ex3)^* mice to generate *Catnb^lox(ex3)/lox(ex3)^;Tg(Col2a1-CreERT2)* animals following several rounds of breeding. The pMes-caBmpr1a mice were generously provided by YiPing Chen (Tulane University, USA), whereas Bmp2^c/c^; Bmp4^c/c^ mice were obtained from Clifford Tabin (Harvard Medical School, USA).

For BMP signaling gain-of-function studies, pMes-caBmpr1a mice were crossed with TgCol2a1-Cre-ERT2 mice to generate *pMes-caBmpr1a;TgCol2a1-Cre-ERT2* animals. For BMP loss-of-function experiments, Bmp2^c/c^; Bmp4^c/c^ mice were bred with *TgCol2a1-Cre-ERT2* mice to obtain *Bmp2^c/c^; Bmp4^c/c^; TgCol2a1-Cre-ERT2* mutants.

### Genotyping

Mice were genotyped using PCR with the following primers:

***TgCol2a1-CreERT2***: forward 5′-CACTGCGGGCTCTACTTCAT-3′, reverse 5′-ACCAGCAGCACTTTTGGAAG-3′.

***Catnb^lox(ex3)/lox(ex3)^*:** forward 5′-GCTGCGTGGACAATGGCTAC-3′, reverse 5′-GCTTTTCTGTCCGGCTCCAT-3′.

***pMes-caBmpr1a***: forward 5′-CGAAGATATGCGTGAGGTTGTGT-3′, reverse 5′-TGGGCGAGAGGGGAAAGAC-3′.

***Bmp2^c/c^:*** forward 5′-GTGTGGTCCACCGCATCAC-3′, reverse 5′-GGCAGACATTGTATCTCTAGG-3′.

***Bmp4^c/c^***: forward 5′-AGACTCTTTAGTGAGCATTTTCAAC-3′, reverse 5′-AGCCCAATTTCCACAACTTC-3′.

### Anterior Cruciate Ligament Transection (ACLT)

We subjected the left hindlimbs of animals to ACLT surgical induction at P70. The surgical procedure was followed as mentioned earlier[24]. In brief, anterior cruciate ligament transection (ACLT) was performed by medially dislocating the patella to expose the femorotibial joint, followed by ACL transection using a microsurgical blade under a stereomicroscope. The patella and quadriceps were repositioned and sutured, and the incision was closed with 5-0 Vicryl sutures. Sham-operated animals underwent joint exposure without ACL transection. Postoperatively, the skin was treated with povidone-iodine, animals received antibiotics for three days, and health was monitored continuously to minimise potential effects on experimental outcomes.

### Wnt-β-catenin gain-of-function experiments

3 doses of tamoxifen (Sigma-Aldrich, 2.5 mg/25g body weight) were administered to *Catnb lox^(ex3)(ex3)^; TgCol2a1::Cre-ERT2* mice at P70 to constitutively activate Wnt-β-catenin signaling in articular chondrocytes.

### Intra-articular delivery of BMP2, LDN-193189, and H_2_O_2_ in the mouse knee joint cavity

To exogenously stimulate BMP signaling, recombinant BMP2 protein (500 ng per dose) was administered unilaterally into the knee joint cavity of 6-month-old mice on two separate occasions, with injections on alternate days. Joint tissues were harvested at 48 hours, 28 days, and 56 days post-injection for molecular and histological analyses. For BMP signaling inhibition, 6 μl of 10 μM LDN-193189 dissolved in 3% 2-(hydroxypropyl)-β-cyclodextrin (w/v in PBS) was administered intra-articularly, while the vehicle control group received 3% 2-(hydroxypropyl)-β-cyclodextrin (w/v in PBS) alone. To induce ROS in adult articular cartilage, two injections (administered on alternate days), each containing 6µl of 2% H2O2 in PBS (v/v), were injected into the joint cavity of 6-month-old C57BL/6J mice. All injections were administered unilaterally into the joint cavity of the mice, except for the LDN-193189 treatment, where the left limb received the drug, and the right knee joint was injected with vehicle. Moreover, we monitored body weight and several other physical parameters to evaluate any potential systemic effects in these animals and found no significant changes in weight or overall health.

### BMP signaling induced NFκB nuclear localisation experiment

BRITER cells were cultured in DMEM supplemented with 10% FBS until passage 2 (P2). Cells were treated with recombinant BMP2 protein (400 ng/mL) for 4 hours, after which they were fixed with 4% paraformaldehyde (PFA). Immunoreactivity against NF-κB was subsequently analysed.

### Articular chondrocytes birth dating experiment

Pregnant C57BL/6J mice at embryonic day 12.5 (E12.5) were injected intraperitoneally with BrdU (100 mg/kg, dissolved in 10 mL PBS). Offspring were harvested at 2 and 6 months of age. BrdU-positive chondrocytes in AC were detected by immunohistochemistry using an anti-BrdU antibody.

### Tissue Processing, histology, and immunohistochemistry

Mice limbs were harvested, and all the tissues were fixed in 4% paraformaldehyde for 16 hours at 4°C. After adequate fixation, the fixed tissues were washed in PBS (phosphate-buffered saline) for 2-3 hours twice to remove any trace amount of PFA. The tissues were then decalcified in 14% EDTA (pH 7.4) for 28 days at room temperature. Then, the tissues were washed in PBS twice for 2 hours for complete removal of EDTA. The tissues were then dehydrated using an alcohol gradient, keeping tissues for at least 4 hours in each ascending alcohol percentage. Dehydrated tissues were embedded and sectioned to obtain 5 µm thick sections. Classical histological analysis was performed using Safranin O/ Fast Green, H&E, and Toluidine Blue staining. For Safranin O/ Fast Green staining, tissue sections were rehydrated and treated with Harris hematoxylin for 5 minutes. Then, it was washed with running tap water and added to 1% Fast Green stain for 5 minutes, dipped in water, then treated with 1% acetic acid, and stained with 0.5% Safranin O for 2 minutes. Then, stained slides were dehydrated using 100% alcohol, transferred to xylene, and mounted in DPX. For immunohistochemistry, tissue sections were initially rehydrated using a descending alcohol gradient. Antigen retrievals were performed specific to the protein molecule to be detected. For collagen detection, acidic 0.05% pepsin was used at 37°C for 30 minutes. Citrate buffer retrieval was the primary choice for all the other antigens, including pSMAD 1/5/9, NFkB, ICAT, BMP2, BMP4, BMPR1A, β-cat, and BrdU. Afterwards, these tissue sections were washed with PBT for 10 minutes three times. 5% goat serum was used to ensure blocking non-specific binding for at least an hour. After blocking, we incubated the tissue sections with primary antibodies against Type II Collagen (DSHB, Cat.no. CIIC1-s, 1:20), Type X Collagen (DSHB, Cat.no. X-AC9-s, 1:20), pSMAD1/5/9 (CST Cat. no.13820S, 1:100), BMP2 (Affinity Bioscience Cat. no. AF5163,1:100), BMP4 (Affinity Bioscience Cat no. AF5175, 1:100), BMPR1A (Invitrogen Cat. no. 38-6000, 1:100), NF-κB (CST, Cat.no.D14E12,1:400) and BrdU (Bio-Rad, Cat.no. AHP2405) overnight at 4°C. Tissue sections were then washed with PBST and incubated with fluorescence-labelled secondary antibodies overnight at 4°C. After antibody incubation, sections were further treated with DAPI (1:1000) for 15 minutes and mounted using the antifade solution. A Leica DM 5000B compound microscope was used to capture and process images.

### Blind Data OARSI Scoring

AC was evaluated using three Safranin O-stained sections from varying depths, following Osteoarthritis Research Society International (OARSI) guidelines (58). The average score of the three sections was used for analysis. Scoring was performed blindly by an expert not involved in the in vivo experiments, using a scale of 0–6, where 0–2 indicates minor osteoarthritic changes and 3–6 indicates significant cartilage erosion.

### Quantification of immunohistochemistry data

The midpoint of the condyle was determined using the meniscus detachment point in longitudinal sections during slide selection. Tissue sections within 50 µm on either side of this midpoint were then used as reference regions for analysis. Both condyles were included in the data analysis; however, all images presented in the manuscript are from the lateral condyle to ensure consistency for the reader. The staining intensity was specifically measured for BMP2 and BMP4, as their expression patterns were dispersed both within the extracellular matrix and the cytoplasm of chondrocytes, making intensity-based quantification more appropriate for these markers. For other markers, such as pSMAD1/5/9, Collagen II, Collagen X, ICAT, and β-catenin, there was a clear distinction between positively and negatively stained articular chondrocytes. As a result, quantification was based on the percentage of positively stained cells.

### 3-D volumetric projections of Micro-computed tomography (µCT) data

Three-dimensional (3D) volumetric projections were generated from µCT scans using CTVol 2.0 software. Scans were performed on an eXplore Locus GE scanner (Micro-CT Facility, Indian Institute of Technology, Kanpur) at 50 kVp and 200 µA, producing images with an initial resolution of 5.86 µm per pixel. Processed images were reconstructed, and 3D iso-surface projections were generated using Parallax Innovations MicroView 2.5.0-RC15 (2.5.0-3557).

### Statistical Analysis

We used GraphPad Prism 8.0.2 software to determine the statistical significance of the OARSI scoring and IHC data. We utilised the Brown-Forsythe and Welch ANOVA, Mann-Whitney test, and unpaired t-test with Welch’s correction to analyse the data’s significance. We carefully considered all relevant statistical assumptions before applying a particular test for data analysis. Specifically, the normality was assessed appropriately, and model validation was conducted using bootstrapping with 10,000 repetitions and a 95% confidence interval. The variance distribution of most data sets was normal and homogeneous, with only minor deviations. As a result, we preferred to use parametric tests rather than non-parametric tests for the statistical analysis. The legends that correspond to the figure provide information about the specific test. The error bars (S.D.) display mean ± standard deviation.

## Supporting information

Revised Main text + Supplementary material

## Acknowledgments

We are extremely grateful to Prof. Yi-Ping Chen (Tulane University, USA) for kindly providing the pMes-caBmpr1a mice. We also thank Prof. Shubha Tole (TIFR, Mumbai, India) for generously sharing mouse strains. We would like to acknowledge Dr Sai Prasad Pydi, Dr Mahima Bharti, Dr Saurav Kumar Jha, Dr Jyoti Tripathi, Ms Archita Mishra, and Ms Anupama for their valuable feedback and constructive criticism during manuscript preparation.

## Author Contributions

AB designed the study and acquired the necessary funds. BK designed the study, prepared the manuscript, and conducted all *in vivo* experiments. S FI completed the OARSI scoring and *in vivo* H_2_O_2_-related experiments. AJ carried out OARSI scoring and BMP 2-induced nuclear localisation of NFkB (*in vitro)*. SA conducted *in vivo* BrdU labelling experiments. ST carried out the OARSI scoring. MMT generated the *Ctnnb1/+* mouse strain. SFI and AJ also contributed to the writing of the manuscript.

## Supplementary Material (SM)

### Supplementary figure legends

**Fig. S1.**
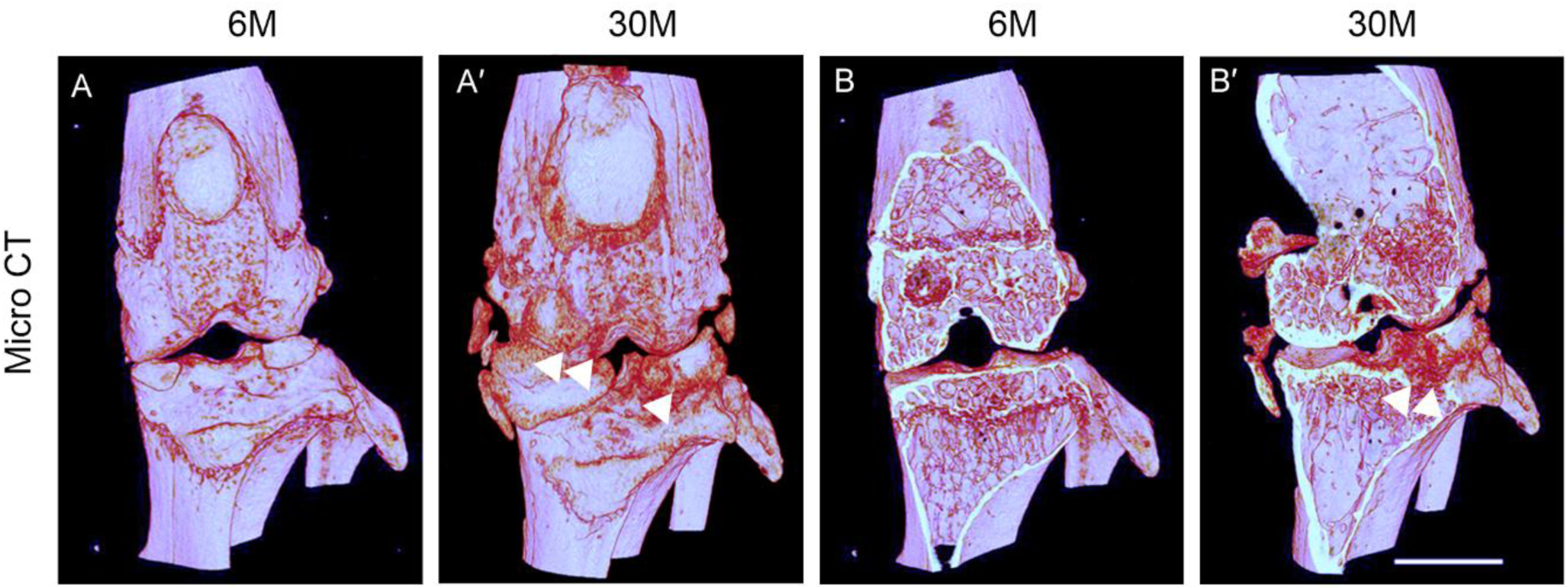
Aging mice’s articular cartilage exhibits radiological changes with aging. (A-A′) 3-D rendering of µCT scan indicates osteophytes, surface roughness, and excessive blood vessels (red colour). (B-B′) Cross-section of the knee joint of 6-month-old (B) and 30-month-old (B′) mice.

**Fig. S2.**
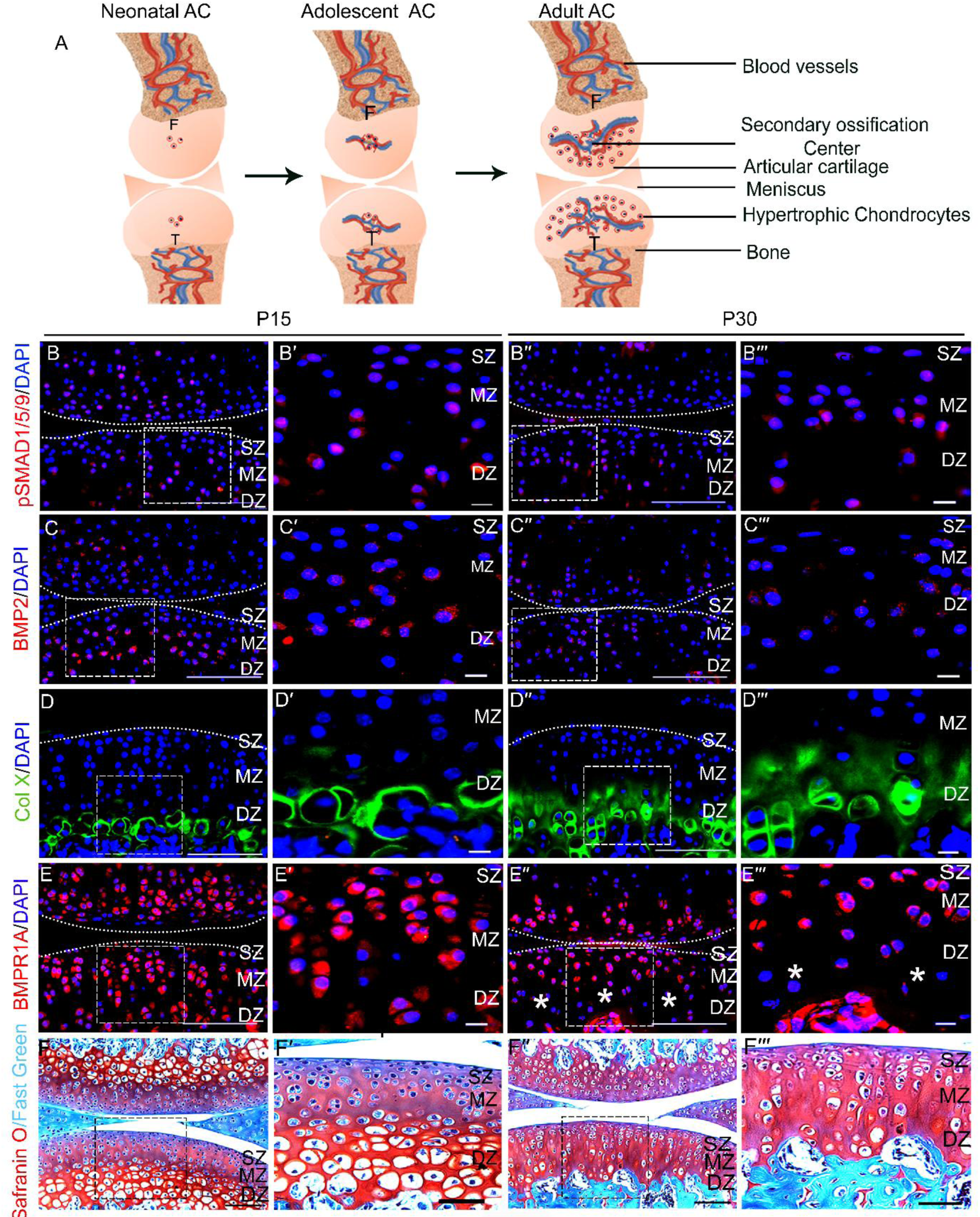
Chondrocyte hypertrophy in the deep zone of developing AC mimics molecular and cellular events of age-associated OA. (A) Schematic representation of the expansion of the hypertrophic domain during the development of postnatal articular cartilage. (B-F′′′) Longitudinal sections through the adult knee joints of C57BL/6J (WT mice) harvested at P15 (B-F′), P30 (B′′-F′′′) of age. Immunoreactivity for pSMAD1/5/9 (B-B′′′), BMP2 (C-C′′′), Col X (D-D′′′), BMPR1A (E-E′′′), Safranin O staining (F-F′′′). B′-F′ are insets for B-F; B ′′′ F′′′ are insets for B′′-F′′ (Inset Scale bar: 20 µm). An asterisk (*) denotes the loss of BMPR1A receptor expression in the deep zone. SZ, MZ, and DZ represent the Superficial, Middle, and Deep zones, respectively. Scale bar: 100 µm; n=5.

**Fig. S3.**
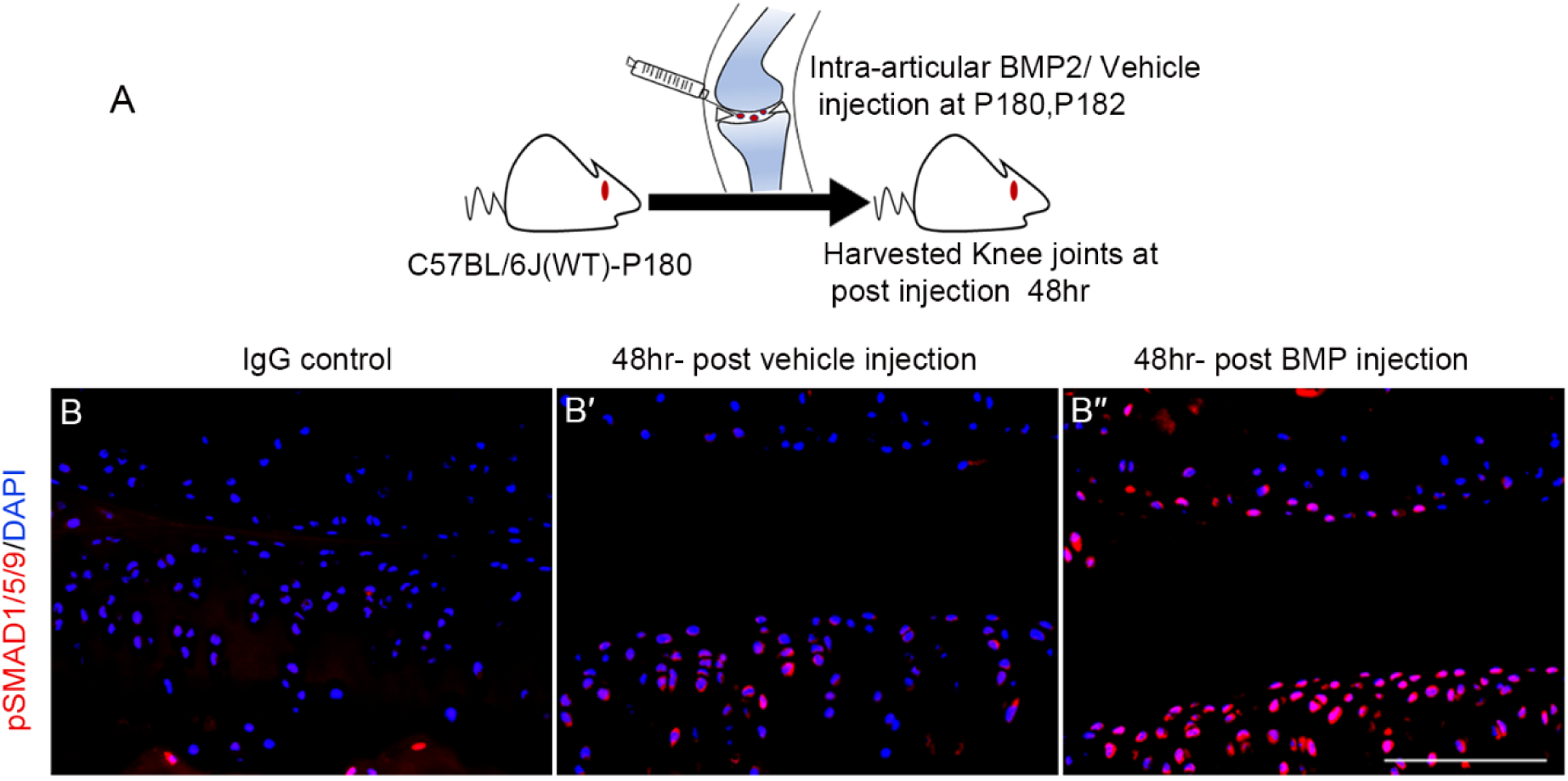
A local supply of BMP2 ligand is sufficient to activate BMP signaling in AC. (A) Schematic for intra-articular injection of BMP2 protein/ Vehicle control at P180, P181 (2 doses) and harvested post-injection 48 hours. (B-B′′) Immunoreactivity for pSMAD1/5/9 on longitudinal sections through the adult knee joints of C57BL/6J (WT mice) of IgG control (B), 48 hours post-vehicle injection(B′), and 48 hours post-BMP2 injection (B′′).

**Fig. S4.**
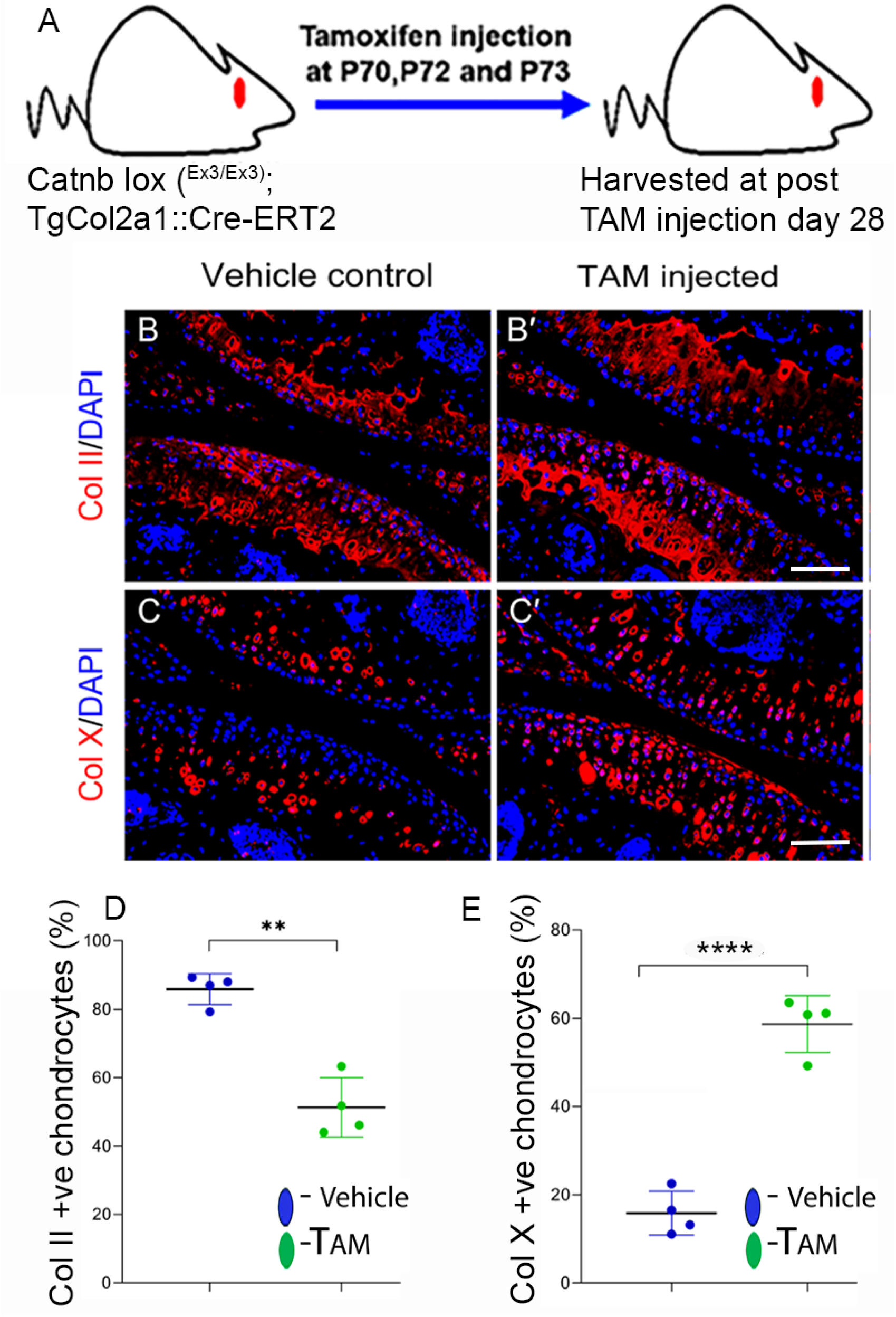
Constitutive active Wnt/β-catenin signaling induces chondrocyte hypertrophy. A) Schematic presentation of activation of Wnt/ β-catenin signaling on tamoxifen injection at P70 of mice. The expression of hypertrophic marker Col X is shown in panel (B-B′), while panel (C-C′) depicts the expression of Col II following tamoxifen injection on Day 28 after injection.

**Fig. S5:**
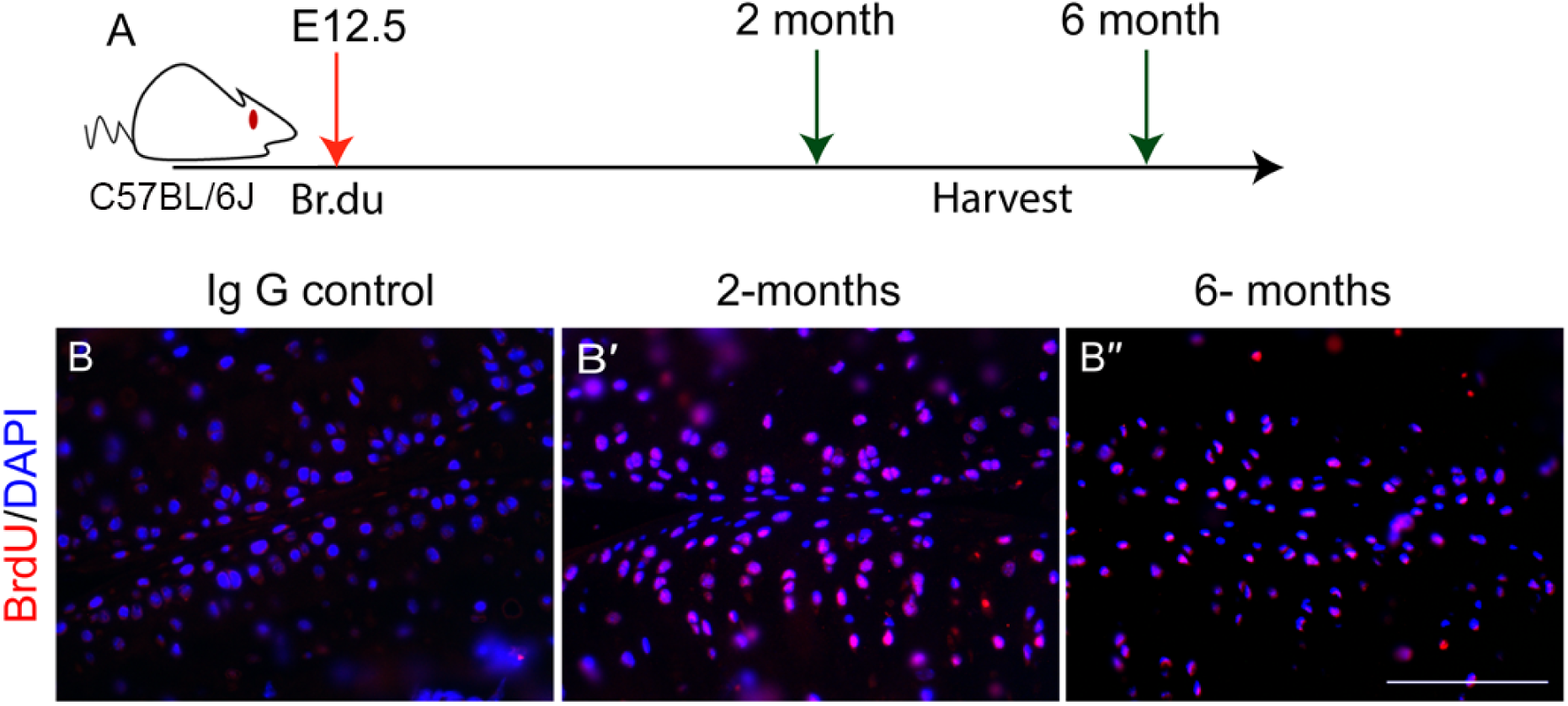
Embryonic articular chondrocytes exist in the adult AC. (A) Schematic showing long-term chase of thymidine analogue BrdU in C57BL/6J mice at E12.5. (B-B′′) BrdU incorporation detected in articular cartilage in IgG control (B) at 2 months (B′) and 6 months (B′′) of age. Scale bar=100 µm; n=3.

**Fig. S6:**
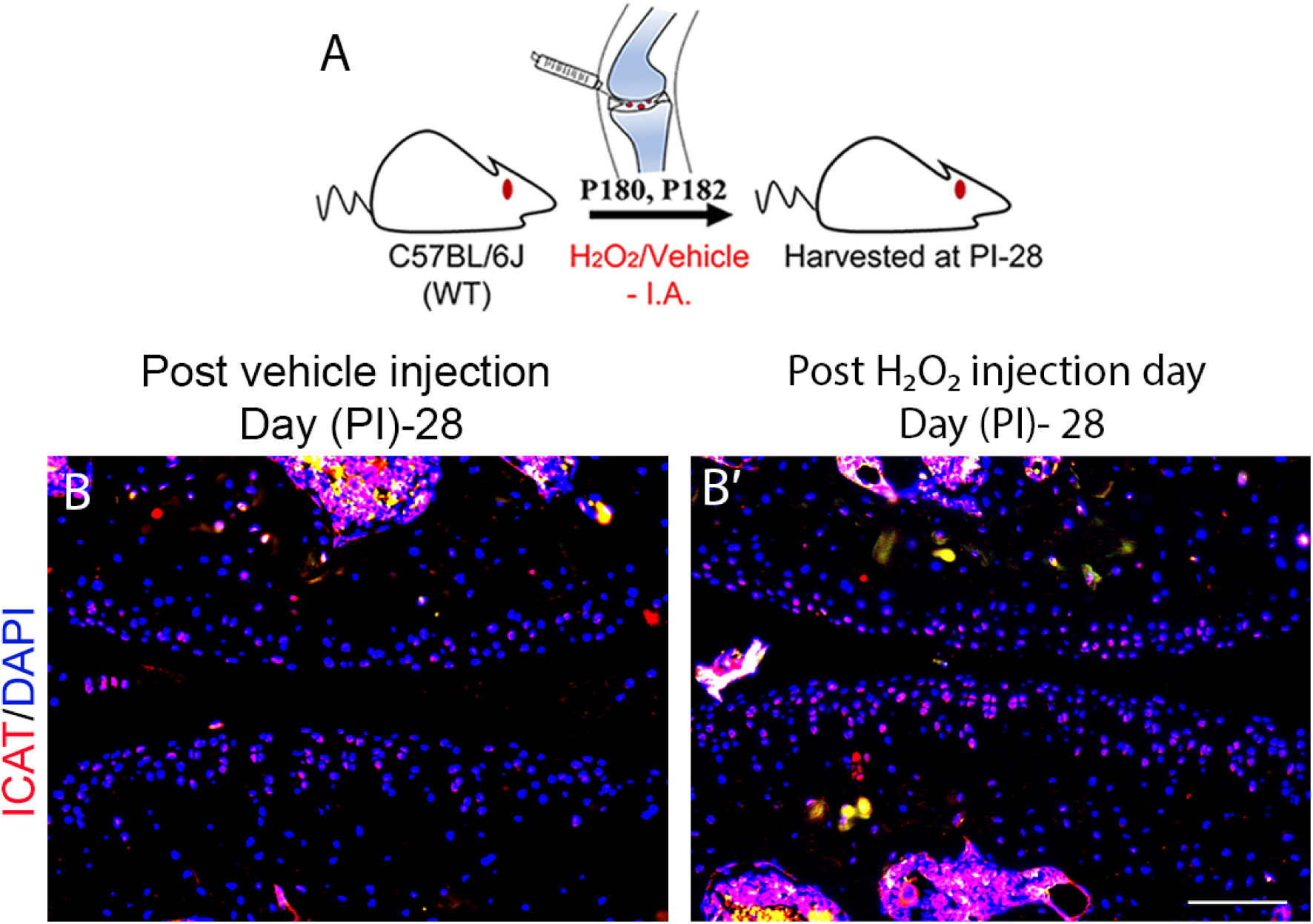
ROS-exposed samples exhibit elevated ICAT expression in articular chondrocytes. (A) Schematic illustrating intra-articular administration of H₂O₂ or vehicle control at P180 and P181 (two doses), with tissue harvested 28 days post-injection. (B–B′) Immunohistochemical staining for ICAT on longitudinal sections of adult knee joints collected 28 days after vehicle treatment (B) and H₂O₂ treatment (B′). Scale bar: 100 µm; n = 5.

