## Supplementary material for "Age-associated spontaneous activation of BMP signaling inhibits the canonical Wnt pathway through ICAT to promote osteoarthritis": Revised Main text + Supplementary material

**Fig. S6: ROS-exposed samples exhibit elevated ICAT expression in articular chondrocytes.**

(A) Schematic illustrating intra-articular administration of $H_2O_2$ or vehicle control at P180 and P181 (two doses), with tissue harvested 28 days post-injection. (B–B′) Immunohistochemical staining for ICAT on longitudinal sections of adult knee joints collected 28 days after vehicle treatment (B) and $H_2O_2$ treatment (B′). Scale bar: 100 μm; n = 5.

### Supplementary figure

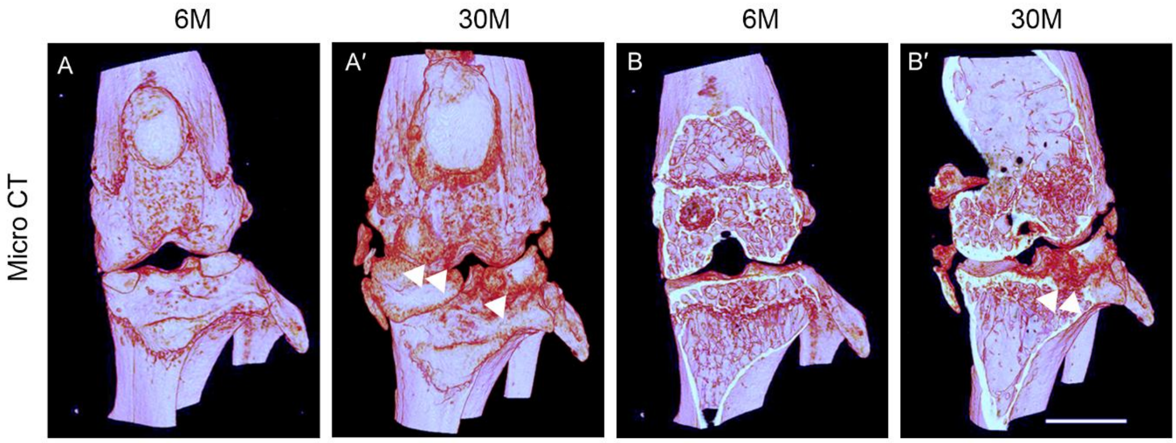


Fig.S1

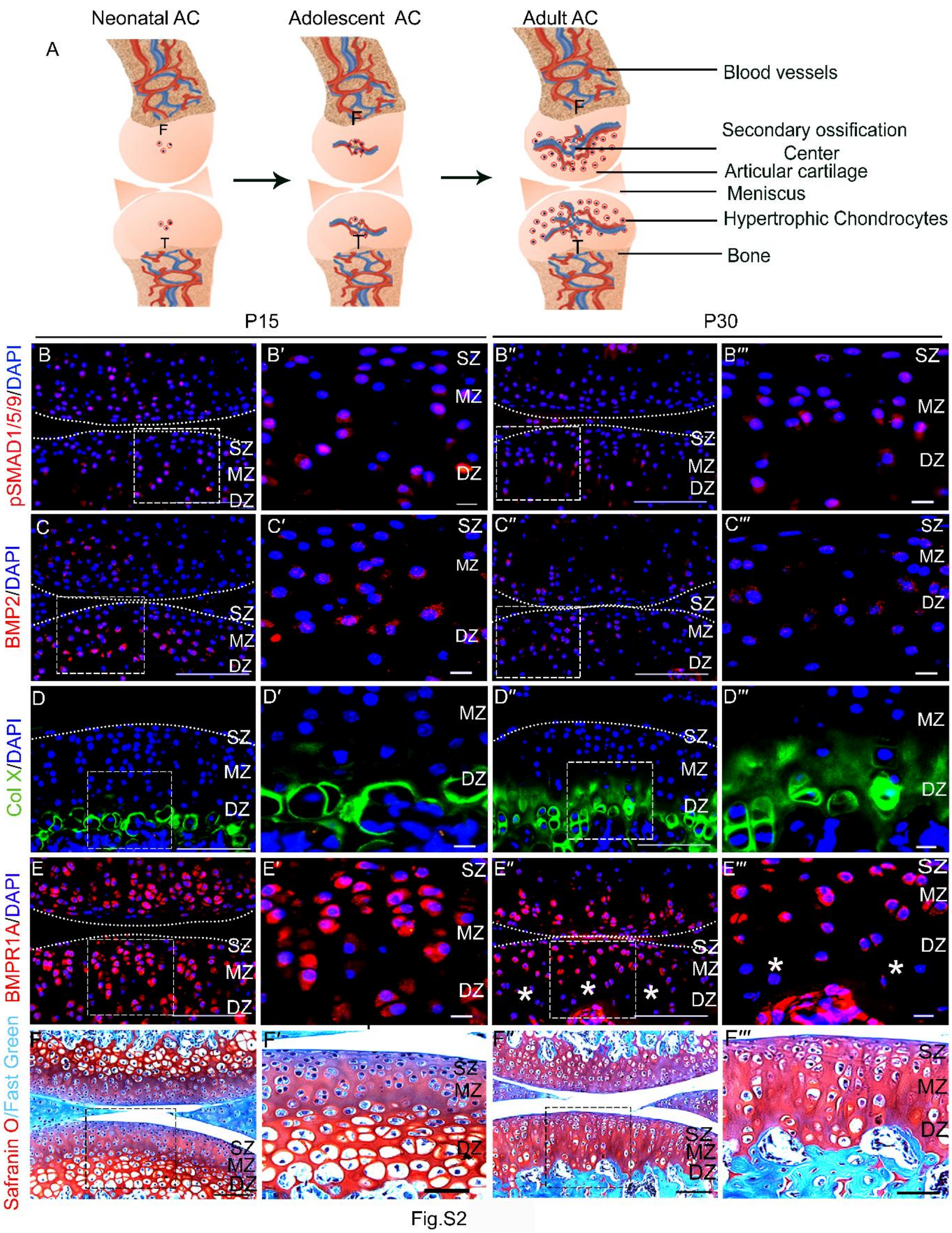

Neonatal AC
Adolescent AC
Adult AC
A
F
T
Blood vessels
Secondary ossification Center
Articular cartilage
Meniscus
Hypertrophic Chondrocytes
Bone
P15
P30
pSMAD1/5/9/DAPI
BMP2/DAPI
Col X/DAPI
BMPR1A/DAPI
Safranin O/Fast Green
SZ
MZ
DZ


Fig.S2

51
52

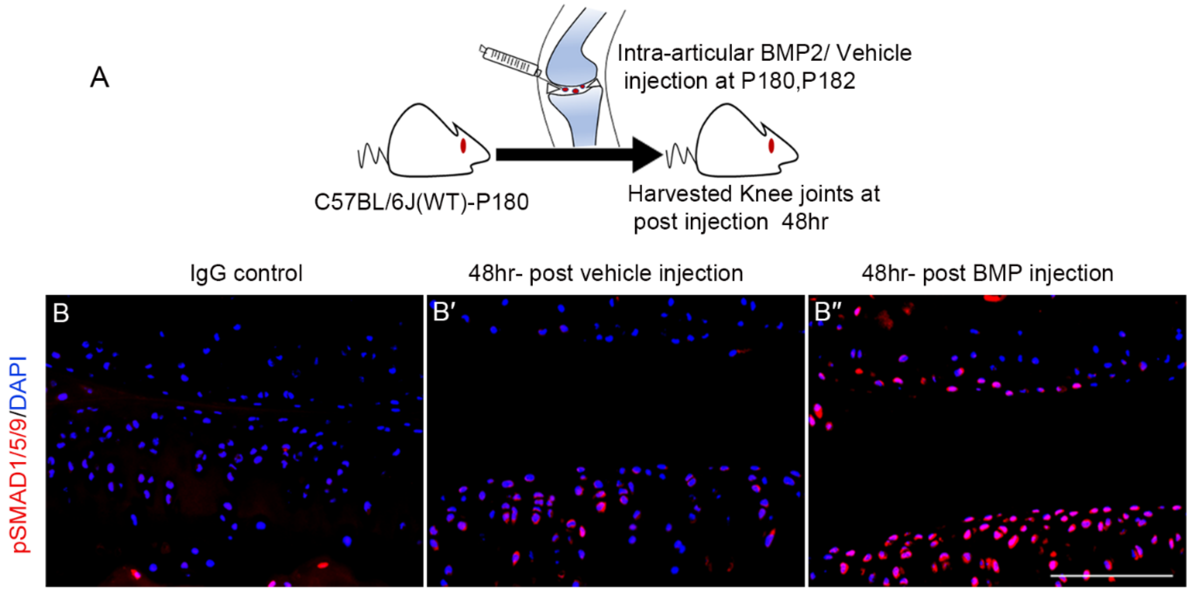
A
Intra-articular BMP2/ Vehicle
injection at P180,P182
C57BL/6J(WT)-P180
Harvested Knee joints at
post injection 48hr
IgG control
48hr- post vehicle injection
48hr- post BMP injection
B
B′
B″
pSMAD1/5/9/DAPI

Fig.S3

53
54

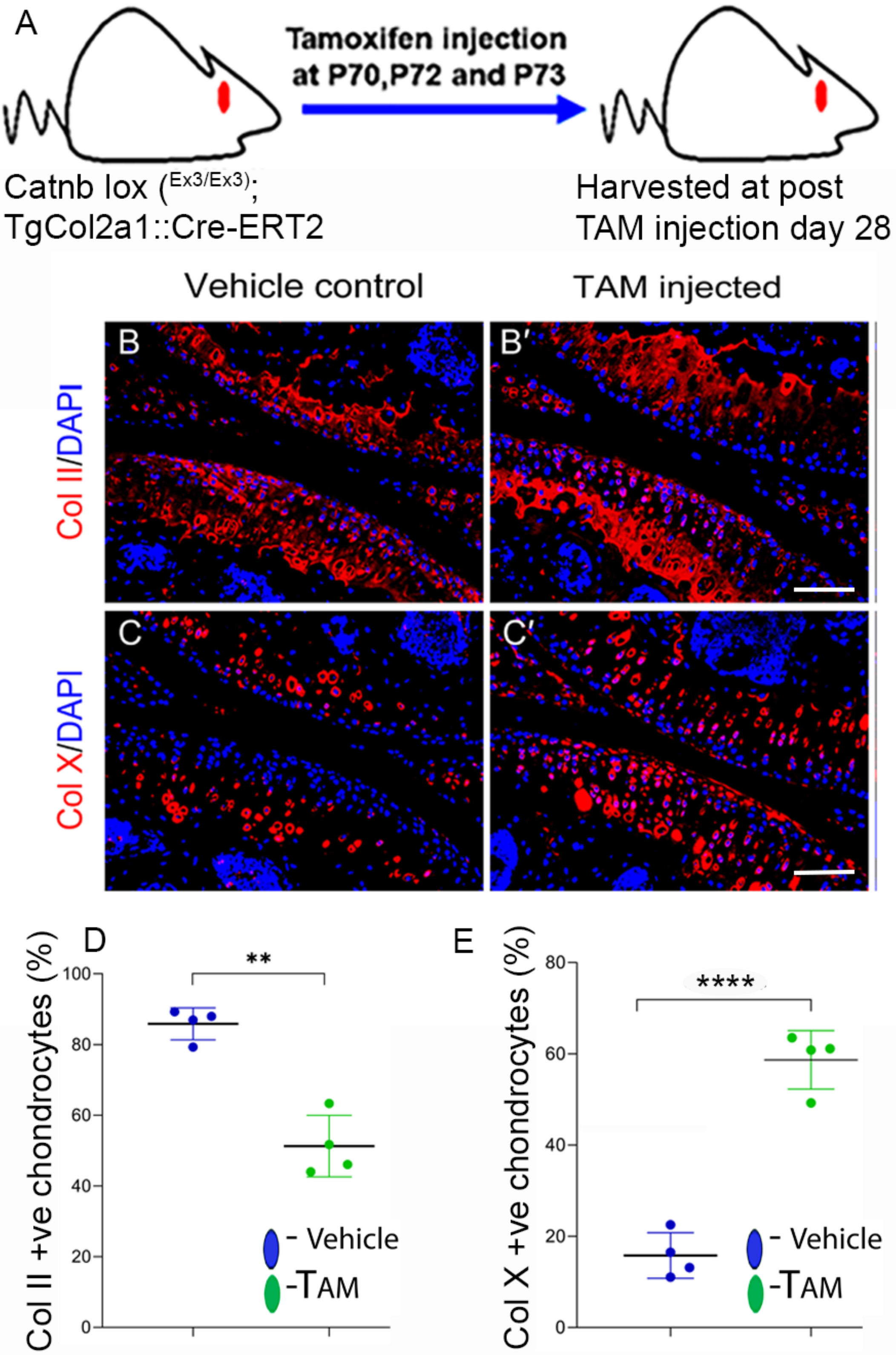
A
Tamoxifen injection at P70,P72 and P73
Catnb lox (Ex3/Ex3);
TgCol2a1::Cre-ERT2
Harvested at post TAM injection day 28
Vehicle control
TAM injected
B
B′
C
C′
Col II/DAPI
Col X/DAPI
D
Col II +ve chondrocytes (%)
**
- Vehicle
-TAM
E
Col X +ve chondrocytes (%)
****
- Vehicle
-TAM

Fig.S4

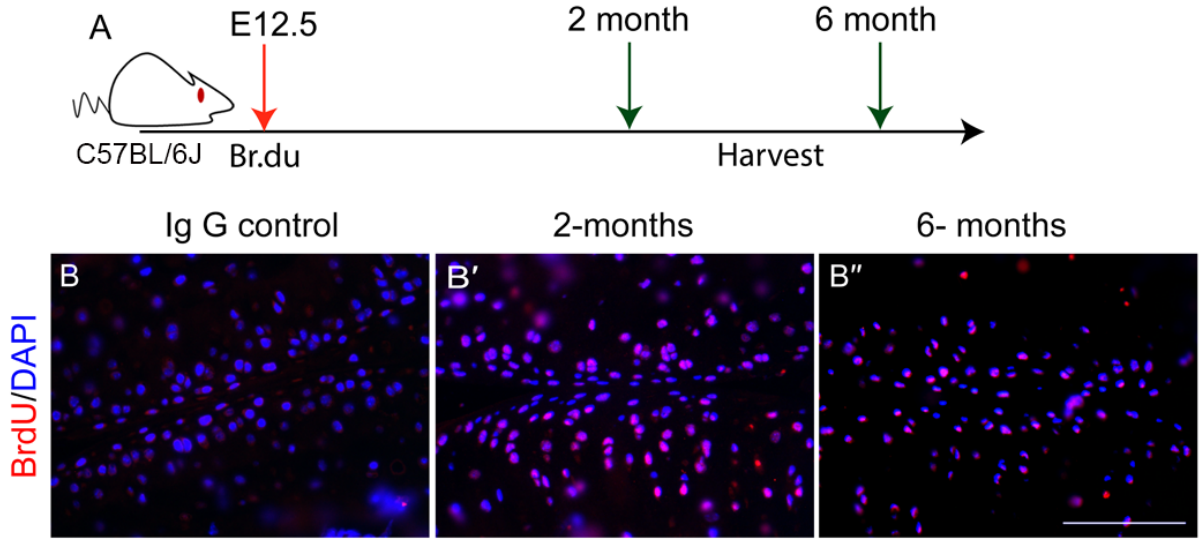
A
E12.5
2 month
6 month
C57BL/6J
Br.du
Harvest
Ig G control
2-months
6- months
B
B′
B″
BrdU/DAPI

Fig.S5

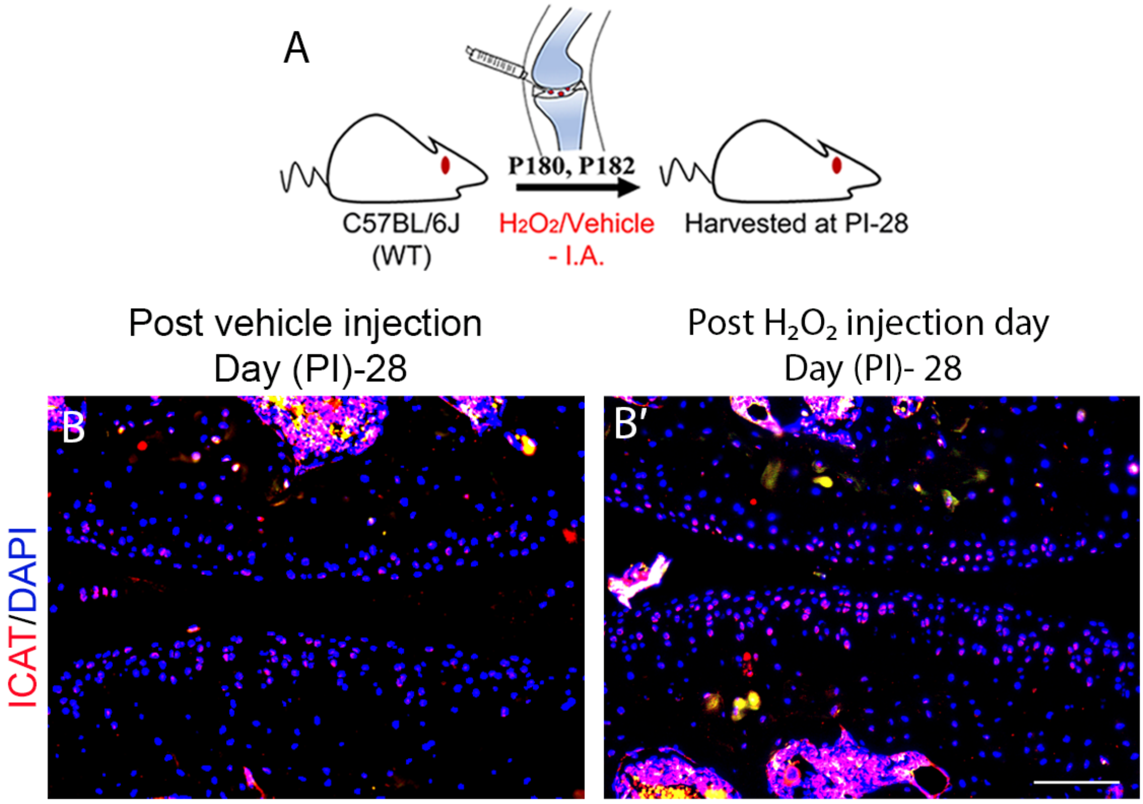
A
P180, P182
C57BL/6J
(WT)
$H_2O_2$/Vehicle
- I.A.
Harvested at PI-28
Post vehicle injection
Day (PI)-28
Post $H_2O_2$ injection day
Day (PI)- 28
B
B′
ICAT/DAPI

Fig.S6
